# Oxidative rearrangement of tryptophan to indole nitrile by a single diiron enzyme

**DOI:** 10.1101/2023.08.03.551874

**Authors:** Sanjoy Adak, Naike Ye, Logan A. Calderone, Rebecca J. B. Schäfer, April L. Lukowski, Maria-Eirini Pandelia, Catherine L. Drennan, Bradley S. Moore

## Abstract

Nitriles are uncommon in nature and are typically constructed from oximes via the oxidative decarboxylation of amino acid substrates or from the derivatization of carboxylic acids. Here we report a third strategy of nitrile biosynthesis featuring the cyanobacterial nitrile synthase AetD. During the biosynthesis of the ‘eagle-killing’ neurotoxin, aetokthonotoxin, AetD converts the alanyl side chain of 5,7-dibromo-L-tryptophan to a nitrile. Employing a combination of structural, biochemical, and biophysical techniques, we characterized AetD as a non-heme diiron enzyme that belongs to the emerging <u>H</u>eme Oxygenase-like <u>D</u>iiron <u>O</u>xidase and Oxygenase (HDO) superfamily. High-resolution crystal structures of AetD together with the identification of catalytically relevant products provide mechanistic insights into how AetD affords this unique transformation that we propose proceeds via an aziridine intermediate. Our work presents a new paradigm for nitrile biogenesis and portrays a substrate binding and metallocofactor assembly mechanism that may be shared among other HDO enzymes.

## Introduction

The nitrile functional group with its short, polarized C–N triple bond is a common feature of medicinal compounds due to its favorable physicochemical and pharmacokinetic properties, metabolic stability, and ability to serve as a bioisostere of carbonyls and halogens.^1,2^ Many nitrile-containing pharmaceuticals are FDA-approved and used for the treatment of several diseases, such as heart failure, hypertension, chronic myeloid leukemia, breast cancer, and fungal infections.^1,2^ Given the importance of nitriles in medicinal chemistry, numerous synthetic methods have been established to install the nitrile functionality.^3^ Nature’s biosynthetic strategies, however, are lesser known as natural nitrile containing compounds are relatively uncommon, and represent approximately 0.1% of natural products. The most common naturally occurring nitriles are cyanogenic glycosides from plants in which they function as defensive agents.^4,5^ Nitriles are also found in arthropods, bacteria, and fungi in which they serve diverse roles as secondary metabolites.^6,7^

The biosynthetic pathways for nitriles are only sparsely described. In plants, N-hydroxylation and subsequent decarboxylation of an amino acid precursor gives rise to an aldoxime intermediate that becomes dehydrated to yield the nitrile (i.e., cyanogenic glycoside), and one such example is the conversion of L-tyrosine (**1**) to (*S*)-dhurrin (**3**) (**Figure 1A, top**). Both aldoxime formation and dehydration are catalyzed by heme-containing enzymes, including dedicated cytochrome P450s.^8 9,10^ In bacteria, the ATP-dependent ToyM is a nitrile-forming enzyme that converts a carboxylic acid to a nitrile via an amide intermediate resulting in the formation of the antibiotic toyocamycin (**6**) (**Figure 1A, middle**).^11^ Although there are proposed gene candidates for nitrile formation in the microbial biosynthesis of the enediyne natural product cyanosporaside^12^ and the macrolide borrelidin^13^, the exact reaction pathways have not been yet delineated.

**Fig 1.**
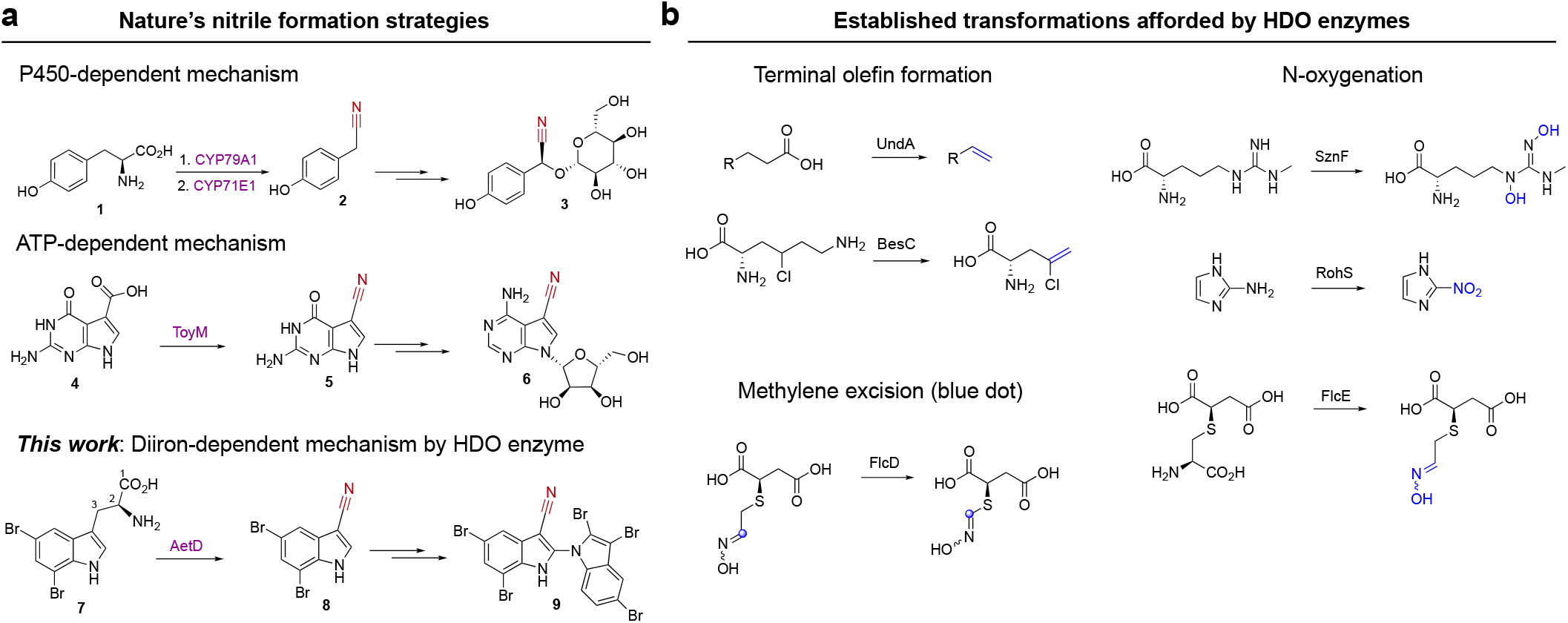
AetD is a member of the HDO enzyme superfamily that catalyzes nitrile formation. **a, Known nitrile biosynthetic enzymes in nature. Top**, Two cytochrome P450s are involved in converting tyrosine (**1**) via an oxime to nitrile **2** during the biosynthesis of the cyanogenic glucoside (*S*)-dhurrin (**3**). **Middle**, ToyM catalyzes the conversion of carboxylic acid **4** via an amide intermediate to nitrile **5** during toyocamycin (**6)** biosynthesis. **Bottom**, AetD converts the alanyl side chain of 5,7-dibromo-L-tryptophan (**7**) to nitrile **8** en route to the biosynthesis of cyanobacterial toxin aetokthonotoxin (**9). b, Previously established reactions of HDO biochemistry**. Nitrile formation expands the known reactivity of HDO enzymes.

Recently, we reported a new nitrile synthase of cyanobacterial origin, AetD, that converts 5,7-dibromo-L-tryptophan (**7**) to 5,7-dibromoindole-3-carbonitrile (**8**) during the biosynthesis of aetokthonotoxin^14^ (**9**) (**Figure 1A, bottom**), a cyanotoxin known for being responsible for the death of eagles.^15^ Although we previously showed that the nitrogen atom from the α-amine of **7** is retained in the nitrile product,^14^ the origin of the nitrile carbon and the fate of the excised carbon atoms were not established. A single enzyme-mediated conversion of tryptophan to indole-3-carbonitrile is unprecedented and offers a new mechanistic paradigm of coupling nitrile formation to carbon deletion biochemistry. AetD shows no homology to ToyM or to cytochrome P450s, even to those associated with formation of the structurally similar indole-3-acetonitrile during the biosynthesis of the plant phytoalexin camalexin.^16^ Instead, AetD utilizes Fe(II) for the conversion of the alanyl side chain of 5,7-dibromo-L-tryptophan to a nitrile.

Our present study dissects the structural and mechanistic features of the AetD-catalyzed reaction through crystallographic analysis of AetD, mechanistic studies employing isotopically labeled substrates, and spectroscopic and kinetic characterization of reaction intermediates. Our work establishes AetD as a new member of the <u>H</u>eme Oxygenase-like <u>D</u>iiron <u>O</u>xidase and Oxygenase (HDO) superfamily^17^, for which chemical transformations include: N-oxygenation (SznF^18^, RohS^19^, FlcE^20^), terminal-olefin formation (UndA^21^, BesC^22^), methylene excision (FlcD^20^) (**Figure 1b**), and the conversion of a protein-derived tyrosine side chain to para-aminobenzoate as part of the biosynthesis of folic acid in *Chlamydia trachomatis* (CADD).^23^ The HDO enzymes can be classified into two subgroups depending on whether the O_2_-reactive complex forms in the absence or in the presence of substrate (i.e., substrate-independent or substrate-triggered, respectively). In substrate-independent HDOs (e.g., SznF)^24,25^ assembly of the diiron cofactor and formation of the O_2_-reactive complex do not require substrate binding. On the other hand in substrate-triggered HDOs (e.g., UndA and BesC)^26-28^, substrate binding likely precedes or facilitates cofactor binding. Here we show that AetD is substrate-triggered and catalyzes a reaction that does not match those previously reported for homologous enzymes, thus expanding the functional repertoire of the HDO superfamily.

## Results

### AetD is a new member of the HDO structural superfamily

A BLAST search of the AetD sequence did not retrieve any significant similarity hits, prompting us to determine the crystal structure of AetD. We first determined the crystal structure of substrate-bound AetD to 2.30 Å resolution by Se-methionine SAD phasing, and subsequently used that structure to solve a higher (2.08 Å) resolution of AetD with substrate-bound and two Fe(II)- and substrate-bound AetD structures to 2.00 Å and 2.30 Å resolution (**Table S1**). AetD appears to be dimeric (**Figure S1**); each AetD protomer has a 7-helical bundle architecture with three core helices that house residues responsible for coordinating the diiron cofactor, and four auxiliary helices (**Figure 2**). Three of the auxiliary (aux) helices aux α1, α3, and α4 pack tightly against aux α2 and the core helices, forming a largely hydrophobic substrate binding pocket (**Figure 2A, B, S2**). This same 7-helical bundle fold was previously observed for HDO enzymes: CADD;^29^ UndA;^21^ BesC;^28^ and SznF^25^ (**Figure S3**). Members of the HDO structural family differ from those of <u>F</u>erritin-like <u>D</u>iiron <u>O</u>xidases/oxygenases (FDOs)^30^ in that they have very labile diiron cofactors, which has posed a challenge for attaining structures of these enzymes with both iron ions bound.^17^ Based on structural comparisons (**Figure S3**), we can now classify AetD as a member of the HDO enzyme superfamily.

**Fig. 2.**
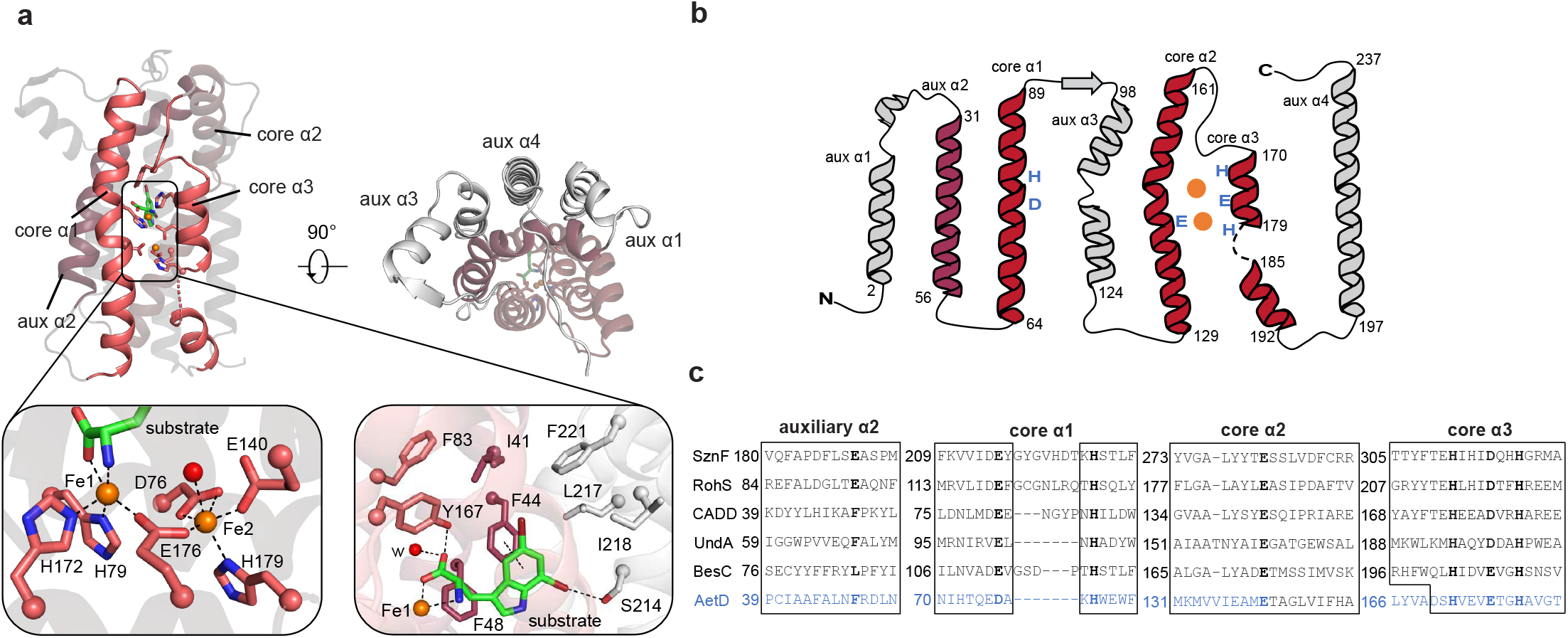
Structure and sequence alignments of substrate- and diiron-bound AetD with other HDO enzymes. **a**, (upper left) Overall 7-helical bundle architecture of AetD. The three core α helices (core α1, α2, and α3) that harbor metal-binding ligands (sticks) are shown in red ribbons. Auxiliary α2 (aux α2) is rendered dark red. The rest of the auxiliary helices are colored in gray. Substrate 5,7-dibromo-L-tryptophan is in sticks (C in green; N in blue; O in red; and Br in brown). (upper right) Top-down view of holo-AetD. (lower left) Diiron (orange spheres) cofactor site with interactions to Fe1 and Fe2 indicated by dashed lines. Water molecule is shown as red sphere. (lower right) Substrate binding site. Substrate is stabilized by hydrophobic interactions, coordination to Fe1, and hydrogen bonds. **b**, Topology diagram showing the architecture of AetD. The six metal-binding residues are labeled in blue. The two orange circles represent the diiron cofactor. **c**, Sequence alignment of AetD with other enzymes in the HDO superfamily. Residues in boxes are part of α helices. The 3-His/3-carboxylate motif (shown in bold) are distributed in three core helices, a characteristic shared among HDO enzymes.

The structure of AetD reconstituted with diiron shows that core helix α1 (nomenclature based on ref 25) provides a monodentate ligand (H79) to Fe1 and a bidentate ligand (D76) to Fe2 (**Figure 2A lower left and 2B**,**C**). Core helix α2 provides a single monodentate E140 ligand to Fe2, whereas core helix α3 presents a characteristic ^170^HX_3_EX_2_H^179^ motif that completes the coordination of the diiron cofactor (**Figure 2A, B, S4**). This metal binding site is similar to that of other HDO enzymes CADD;^23,29^ UndA;^21,26,31^ BesC;^22,27,28^ and SznF^18,24,25^ (**Figure S3, S4**).

Interestingly, in these structures, the core helices often deviate from classical helical structures in order to appropriately position the metal-binding ligands. For example, α1 helix in AetD, UndA^26^ and BesC^28^ has a kink at its halfway point where the helix contributes metal-binding ligands into the active site (**Figure S3D and S3E**); in CADD^29^ and SznF^25^, α1 is completely interrupted by a sizeable loop at this same location (**Figure S3C and S2F**). Core helix α3 is also unusual in that its C-terminus is highly flexible allowing the helix to unwind (**Figure S3, S4**), presumably to provide access to substrates and/or cofactor binding sites.^21,25,28^

Structural comparisons suggest differences in α1-3 and aux α2 that are likely important for substrate specificity. For example, α3 starts later in AetD than in other HDO enzymes (**Figure 2C**), resulting in the placement of α3 residue Y167 directly into the active site where it can hydrogen bond to the substrate carboxylate oxygen (**Figure 2A, C, S3**). Whereas aux α2 helices in SznF and RohS contribute Fe1 ligand E189 and E93, respectively (**Figure 2C, S3**), this residue is F48 in AetD and its sidechain stacks against AetD’s substrate (**Figure 2, S3**). The absence of a Glu ligand in AetD provides an open coordination site on Fe1 for ligation of the substrate carboxylate and amine nitrogen (**Figure 2A**).This direct interaction between the polar end of the substrate and Fe1 has also been observed in the substrate-bound structure of the HDO enzyme UndA^26^ (**Figure S5**). In AetD, the substrate is also anchored in the active site by hydrophobic interactions, including a putative π-π interaction with F44 (centroid-to-centroid distance of 4.0 Å), and a 3.0 Å O-H···Br hydrogen bond between S214 and the substrate’s Br substituent (**Figure 2A lower right, S2**).

### Conformational changes of core helix α3 in AetD accompany cofactor assembly

The series of crystallographic snapshots obtained here have allowed us to evaluate conformational rearrangements associated with diiron cofactor assembly. Despite much effort, we were unable to obtain a structure of AetD with iron-bound in the absence of substrate. However, we were able to obtain a substrate-bound co-crystal structure of AetD at 2.08 Å resolution (see **Table S1**), showing excellent omit map density for 5,7-dibromo-L-tryptophan in the absence of iron (**Figure 3A, S2**). In this structure, the carboxylate and amino nitrogen of substrate, which coordinate Fe1 when iron is present, now hydrogen bond to Fe1 ligands H79 and H172, pre-organizing these residues for Fe1 coordination (**Figure 3B**). Thus, the three ligands that exclusively coordinate Fe1 (the amino acid moiety of substrate, H79, and H172) are all positioned for Fe1 binding when substrate is present. Diiron bridging residue E176, and Fe2 ligand H179, on the other hand, are part of a highly flexible region of core helix α3 (E176 to T183) and are disordered (**Figure 3A and S6A**). Of the residues that bind Fe2 (D76, E140, E176, and H179), only D76 of core helix α1 has good omit electron density (**Figure 3A**). Thus, substrate binding appears to pre-organize AetD to bind Fe1 but not Fe2.

**Fig. 3.**
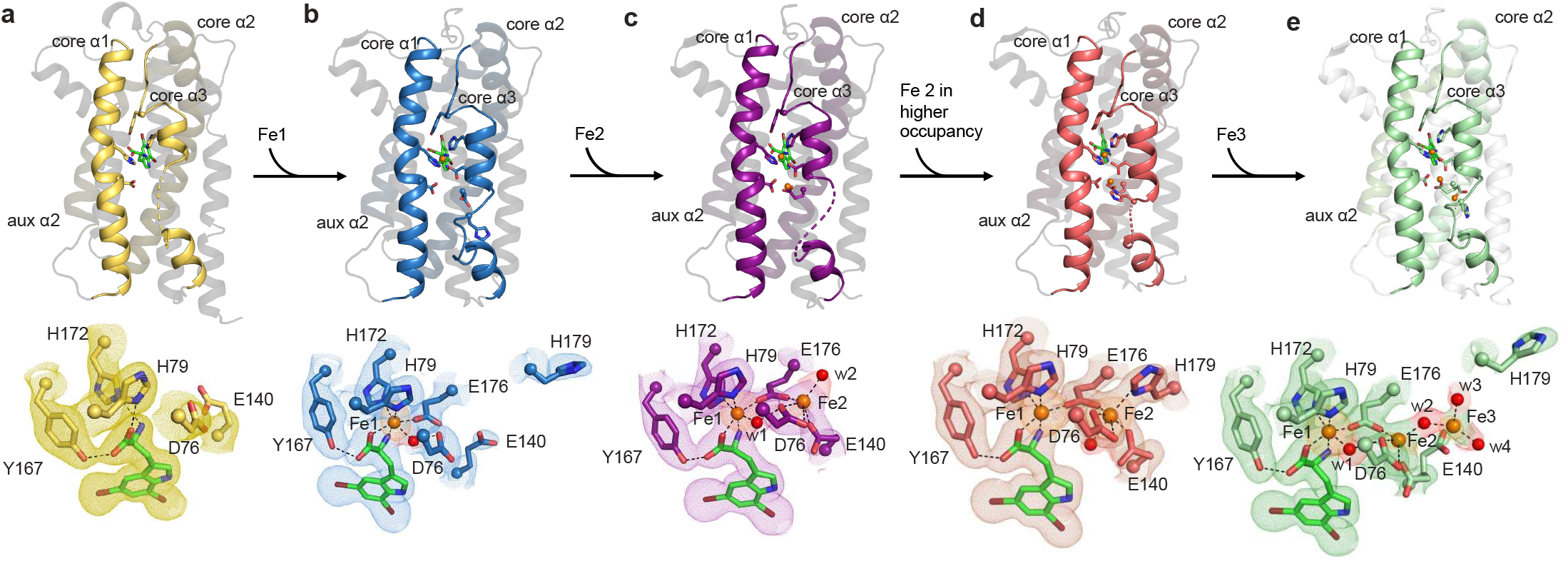
Crystallographic snapshots of substrate binding and diiron cofactor assembly. **a**, Substrate-bound-only, pre-cofactor-assembly state. The amino acid moiety of the 5,7-dibromo-L-tryptophan substrate (green carbons) forms hydrogen bonds with Y167 and Fe1 (orange sphere) ligands H172 and H79. Core helix α3 only has one ordered helical turn. **b**, Substrate and Fe1 bound, cofactor partially assembled state. **c**, Substrate bound and cofactor assembled, Fe2 in low occupancy state. **d**, Substrate and both Fe bound, cofactor fully assembled state. **e**, Substrate bound and cofactor assembled, with a third iron bound at the cofactor. In all structures, substrates form similar interactions with the protein (omitted for simplicity). Selected amino acid side chains are shown in colored sticks, with alpha carbons of these residues in spheres. Water molecules are shown as red spheres. 2mF_o_-DF_c_ composite omit maps contoured at 1.0σ are shown in meshes that match individual color schemes of the enzyme. Iron anomalous difference maps are contoured at 3.0σ and are shown in orange meshes. Omit electron density of selected water molecules are shown in red meshes.

We obtained a snapshot of AetD with one iron bound (Fe1), by expressing the protein in minimal media with Fe(II) supplementation and co-crystallization of the purified AetD with substrate in the absence of additional Fe(II). This substrate-bound structure with a partially-reconstituted-cofactor was determined to 2.3 Å-resolution structure with two protomers in the asymmetric unit (**Table S1**). One protomer (chain A) has a single iron bound at high occupancy in the Fe1 site based on an iron anomalous difference map (**Figure 3B**). The other protomer (chain B), described below, appears to have partial occupancy at both the Fe1 and Fe2 sites based on the iron anomalous difference map (**Figure 3C**). First, for chain A, the single iron (Fe1) is coordinated by H79, H172, E176 and substrate as described above. D76 is positioned similarly to the substrate-alone structure, but now makes a hydrogen bond with a water molecule that is additionally coordinated to Fe1, which attains an overall octahedral geometry. Core helix α3 is more ordered compared to the pre-cofactor assembly state (i.e., substrate-bound-only state, **Figure 3A**); there are two additional helical turns in the segment D170 to T177; T177 to I186 is now ordered as a non-helical loop (**Figure S6A, B**), and the Fe-bridging ligand, E176, and the Fe2-binding ligand, H179, are ordered adjacent to the Fe2 binding site (**Figure 3B**). The density is improved for the Fe2 ligand E140 (**Figure 3B**). The net result of these changes is that D76, E140, and E176 are now available to interact with an incoming Fe2.

Chain B of the partially reconstituted active site structure harbors two Fe ions. Fe1 has a similar high occupancy to Fe1 in chain A (∼84%) with the same ligands and coordination geometry. Fe2 has approximately half of the occupancy of Fe1 (∼43%), which is why we are referring to this structure as partially reconstituted. Fe2 attains a square pyramidal geometry, coordinated by E176, D76, E140, and a water molecule (w2). The only missing ligand is H179, which is in the disordered segment (T177 to I186) of core α3 (**Figure 3C**).

In an attempt to obtain a fully-reconstituted diiron-bound structure of AetD, we took crystals grown under the conditions used for the partially-reconstituted AetD structure and soaked them in a large excess (20 mM) of Fe(II). We obtained a 2.0 Å-resolution structure, again with two protomers in the asymmetric unit (**Table S1**). Based on iron anomalous difference maps, one protomer (chain B) has two Fe ions bound and appears to represent the fully-reconstituted diiron cofactor (**Figure 3D**). The other protomer (chain A) has three Fe ions bound (**Figure 3E**). Presumably the third Fe is an artifact of the high Fe(II) concentrations used in the crystal soaking.

In chain B with the fully-reconstituted diiron cofactor, the occupancy of Fe1 (92%) is still higher than for Fe2 (63%), but Fe2 now is now fully coordinated (**Figure 3D**). For the first time, H179 shows well-defined electron density and is clearly coordinated to Fe2. The rest of the coordination sphere for Fe2 is completed by the bridging carboxylate E176, the bidentate ligand D76, the monodentate ligand E140, as well as a water molecule *trans* to H179, resulting in an octahedral geometry for Fe2. Overall, the metal-binding region of core helix α3 (D170 to H179) adopts the most ordered backbone conformation (3 helical turns) among all our structures (**Figure S6C**). Beyond H179, however, the flexible region of core helix α3 is disordered. In chain A, the presence of a third Fe(II) disrupts the Fe2 coordination environment. The occupancy of Fe2 drops to ∼26% and Fe2 has fewer protein ligands (**Figure 3E**). Fe2 ligand E140 shows alternate conformations, one ligating Fe2 and one ligating Fe3, while H179 flips away and no longer coordinates any of the Fe ions. Fe3 is also at low occupancy (∼19%) and is coordinated by three water molecules (w2, w3, and w4) in addition to being coordinated by one conformation of E140 (**Figure 3E**). Fe1 occupancy and coordination geometry is unchanged (**Figure 3E**). The Fe3 position is close to the bulk solvent and may represent an entry position for iron ions to access the cofactor-binding site.

### Source of the carbon atom in the nitrile functional group

With crystallographic snapshots suggesting that substrate binding precedes the assembly of the diiron cofactor, our next step in delineating the mechanism of AetD was to probe the individual fates of the alanyl side chain carbon atoms of the substrate 5,7-dibromo-L-tryptophan (**7**) to identify the carbon that is the origin of the nitrile group in **8**. Employing cell lysates of *Escherichia coli* expressing the *Salmonella enterica* tryptophan synthase (TrpS),^32^ we enzymatically synthesized three ^13^C-labeled isotopologs of tryptophan: [2-^13^C_1_]-L-tryptophan, [3-^13^C_1_]-L-tryptophan, and [1,2,3-^13^C_3_]-L-tryptophan with indole and labeled L-serine as the two building blocks. The single-component flavin-dependent halogenase AetF was then used to convert the ^13^C-labeled L-tryptophan moiety into the corresponding ^13^C-labeled substrate **7** for AetD.^14^

LC-MS analysis of the reaction of AetD with the ^13^C-labeled 5,7-dibromo-L-tryptophans showed a 1 Da mass increase in the nitrile product only in the presence of [3-^13^C_1_]-5,7-dibromo-L-tryptophan and [1,2,3-^13^C_3_]-5,7-dibromo-L-tryptophan, establishing that C3 is retained in the nitrile product, while C1 and C2 are released (**Figure 4A and S7**). This result implies that the nitrogen atom attached to C2 must migrate to C3 to become the nitrile group, revealing an unusually remarkable rearrangement reaction.

**Fig. 4.**
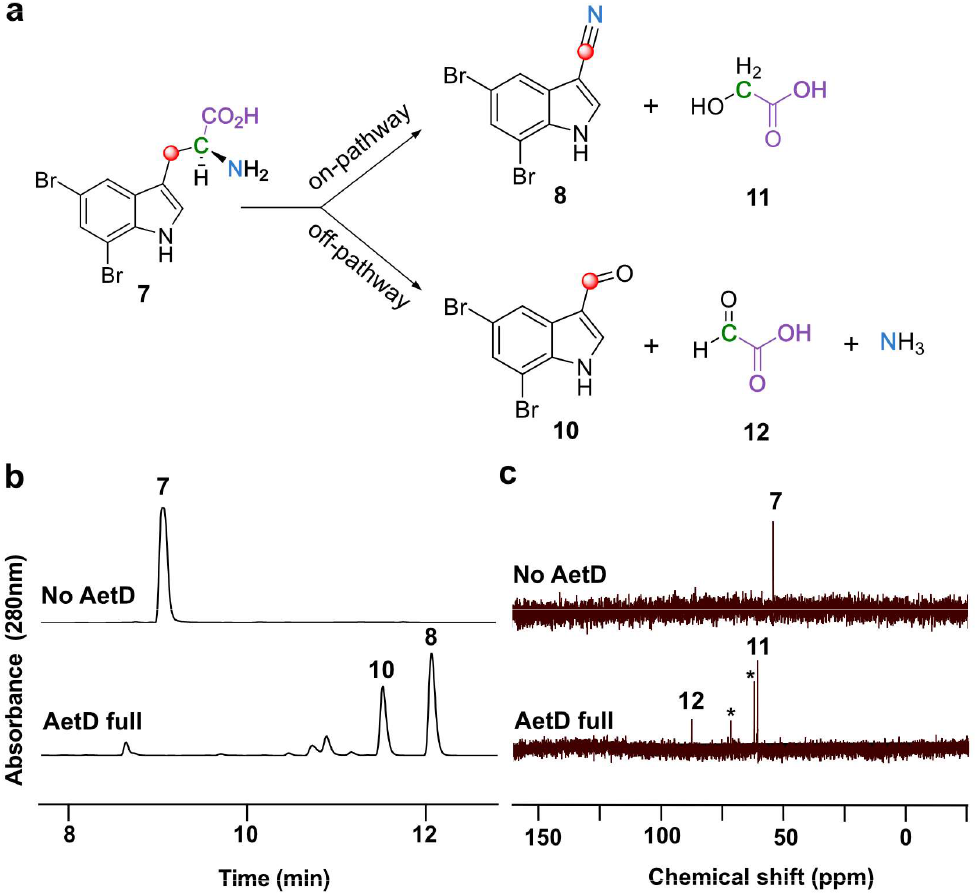
Characterization of all products in the AetD catalyzed reaction. **a**, Schematic showing the product distribution of AetD catalyzed reaction in native and shunt pathways. Native pathway yields 5,7-dibromoindole-3-carbonitrile (**8**) and glycolic acid (**11**) as products, while 5,7-dibromoindole-3-carbaldehyde (**10**), glyoxylic acid (**12**), and ammonia are the products in the shunt pathway. Labeling experiments with ^13^C-labeled isotopologues of the substrate confirmed the fate of the carbon atoms of the alanyl side chain and accordingly color-coded. See Fig. S7–S12 for full set of LC chromatograms and MS and NMR spectra. **b**, LC chromatogram showing the formation of shunt product **10** in addition to native **8** in the AetD full reaction assay. **c**, ^13^C-NMR spectrum of AetD catalyzed reaction in the presence of [2-^13^C]-5,7-dibromo-L-tryptophan (**7**). In the full reaction assay, the C-2 peak corresponding to the substrate **7** disappeared, while two new ^13^C peaks appeared which we characterized as the alcohol and aldehyde carbons of **11** and **12**, respectively. * glycerol peak from the AetD protein stock.

### 5,7-dibromo indole-3-carbaldehyde is a shunt product

HPLC analysis of the AetD reaction with 5,7-dibromo-L-tryptophan (**7**), however, showed the presence of two product peaks (**Figure 4B**). In addition to previously reported 5,7-dibromoindole-3-carbonitrile (**8**), we established 5,7-dibromo-indole-3-carbaldehyde (**10**) as a previously unreported second product. This assignment was confirmed by high-resolution mass data and comparison to chemical standards (**Figure S8**). We incubated **10** in the presence of AetD along with ammonium salt or glycine to explore whether it could be further processed to the nitrile. Under these conditions, however, we did not observe any further transformation, suggesting that **10** is a shunt product and not an on-pathway intermediate. ^13^C-Labeling experiments revealed that C3 is retained in the aldehyde product like the native nitrile product (**Figure S9**). Although no previous biosynthetic routes exist for compound **10**, the des-bromo version of the compound has been previously reported.^33-35^ To note, indole-3-carbaldehyde is produced by plants^33^ as well as human gastrointestinal microbiota^34,35^ and is proposed to be biosynthesized from tryptophan in a multi-enzyme-catalyzed reaction. A more structurally similar compound, 6-bromoindole-3-carbaldehyde, was reported from an *Acinetobacter* sp. bacterium associated with the ascidian *Stomozoa murrayi*.^*36*^ However, the biosynthesis of this compound has not yet been reported.

### The fate of the released two carbon fragments corresponding to nitrile and aldehyde product

Having established the source of the carbon atom in both nitrile and aldehyde products **8** and **10**, we next explored the fate of the excised carbon atoms. Reaction of AetD with [2-^13^C_1_]-5,7-dibromo-L-tryptophan yielded two labeled product peaks in the ^13^C NMR spectrum, one at 61 ppm and the other at 88 ppm (**Figure 4C**). These two peaks were assigned to glycolic acid (**11**) and glyoxylic acid (**12**), respectively, based on their chemical shift values and comparison with standards (**Figure S10**). This assignment was corroborated by two additional experiments. Reaction of AetD with [1,2,3-^13^C_3_]-5,7-dibromo-L-tryptophan gave rise to doubly labeled glycolic and glyoxylic acids, resulting in the additional splitting of the ^13^C NMR peaks due to ^13^C-^13^C J-couplings, thereby demonstrating that both carbon atoms originate from the same substrate molecule (**Figure S11**). The identity of the glyoxylic acid co-product was independently confirmed by *o*-phenylenediamine derivatization followed by MS analysis (**Figure S12**). Comparison of the relative peak intensities in the ^13^C-NMR and LC-MS spectra showed that **11** is the on-pathway side product, whereas **12** is a shunt pathway side product.

### Accumulation of μ-peroxodiiron (III) species in the AetD reaction

Because all known members of the diiron HDO superfamily carry out their oxidative reactions employing μ-peroxodiiron (III) species,^24,26-28^ we sought to observe any such intermediates in AetD. Reaction of an anoxic solution of Fe(II)-reconstituted AetD with O_2_-saturated buffer results in the slow development (k_obs_∼ 0.06 s^-1^) of an absorbance feature with λ_max_ ∼ 325 nm that signifies oxidation of Fe(II) to Fe(III) (**Figure 5A**). By contrast, when 5,7-dibromo-L-tryptophan is added to the ferrous containing solution of AetD and then reacted with O_2_, a new spectrum with absorption maxima at 350 nm and 625 nm rapidly develops (k_obs_ ∼ 8.2 s^-1^) (**Figure 5B**) and exhibits a dependence on the O_2_ concentration (k[O_2_] = 1.12 10^4^ M^-1^ s^-1^) (**Figure S13**). The absorption features of this species resemble those of μ-peroxodiiron(III) intermediates in non-heme diiron oxygenases and oxidases (i.e., FDOs and HDOs) and are assigned to peroxo-to-Fe(III) charge transfer transitions.^37^ The 625-nm absorbing complex is transient and decays with an observed rate constant k_obs_ = 0.02 s^-1^ (**Figure 5C**). The substrate-triggered accumulation of this species and its transient nature suggest that it may represent an intermediate in the reaction to yield the nitrile compound. Interestingly, we observed a second species with a λ_max_ ∼ 510 nm forming with a slower observed rate constant (k_obs_ = 7 s^-1^), accumulation of which, however, is also dependent on the concentration of O_2_ (k[O_2_] = 10^4^ M^-1^ cm^-1^) (**Figure S13**). The absorption features of this second species could also be consistent with a μ-peroxodiiron(III) species for which a range of λ_max_ have been reported to be within 450 – 700 nm.^37^ Decay of the 625-nm absorbing complex is not O_2_-dependent, unlike the formation of the 510-nm absorbing species, suggesting that these two species represent separate events upon reaction of ferrous complexes with O_2_ that perhaps differ in the degree of substrate oxidation. The kinetics of the O_2_-dependence of the 625-nm species corroborates that no prior AetD-O_2_ adducts accumulate to a detectable level in the reaction. The 510-nm complex decays with accumulation of a UV-visible species at 350 nm that resembles that of μ-(hydr)oxodiiron(III) end reaction complexes.

**Fig. 5.**
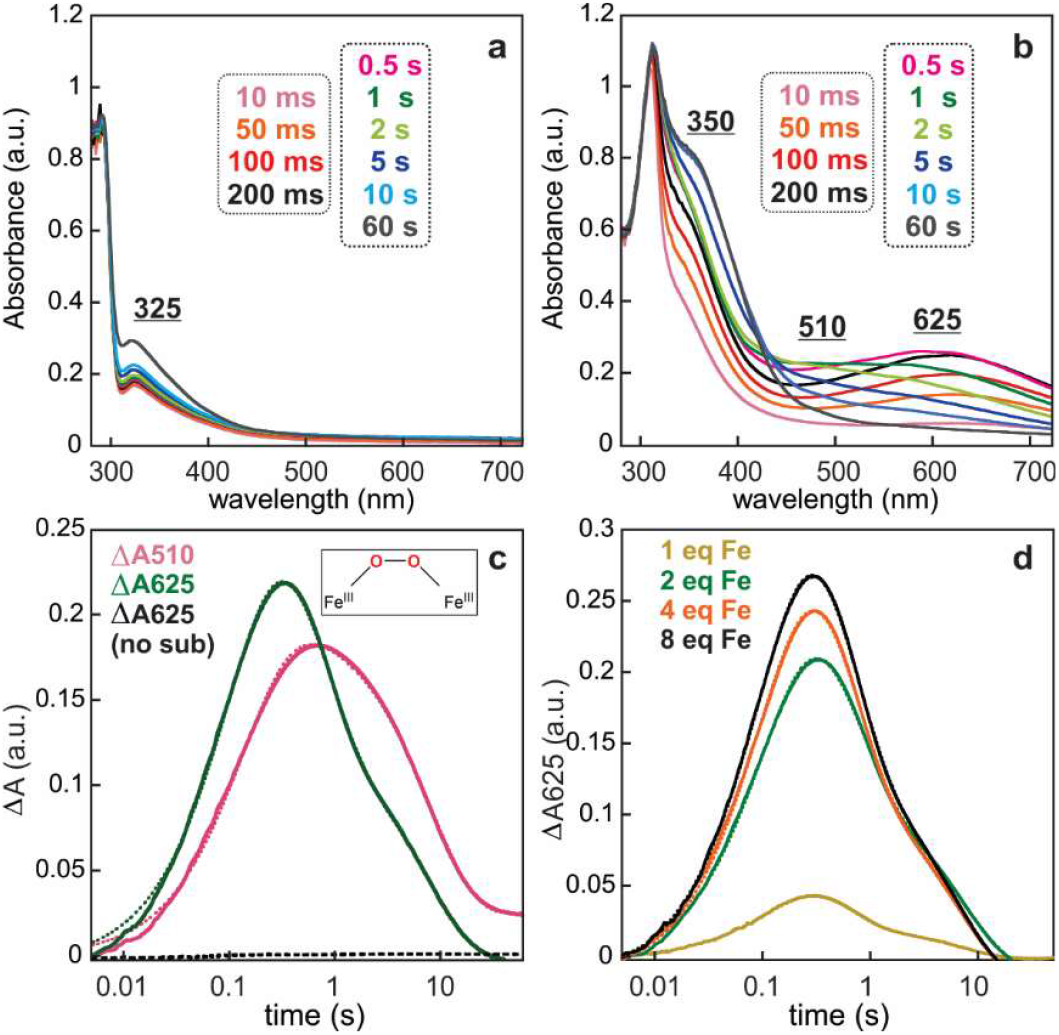
SF-Abs spectra that demonstrate accumulation of intermediate(s) in the reaction of the Fe(II)·AetD complex with O_2_ only in the presence of 5,7-dibromo-L-tryptophan at 5 °C. Absorption spectra acquired after rapid mixing of an O_2_-free solution of AetD (0.30 mM) and Fe(II) (0.60 mM, 2 molar eq) in the (**a**) absence or (**b**) presence of 2 mM 5,7-dibromo-L-tryptophan with an equal volume of O_2_-saturated buffer (1.8 mM). **c**, Kinetic traces showing the accumulation and decay of the absorbing intermediates as a function of time in the presence of 2 mM 5,7-dibromo-L-tryptophan (initial concentration). The control lacking the substrate is shown in black. **d**, Kinetic traces of the intermediate with absorption maxima λ = 625 nm as a function of 1 molar eq Fe(II) (gold trace), 2 molar eq Fe(II) (green trace), 4 molar eq Fe(II) (orange trace), and 8 molar eq Fe(II) (black trace) in the presence of 2 mM 5,7-dibromo-L-tryptophan (initial concentration).

The kinetics of formation of the 625-nm and 510-nm transient species in AetD exhibit a similar dependence on Fe(II) as previously observed for the μ-peroxodiiron(III) intermediate in BesC.^27,28^ Addition of Fe(II) in excess stoichiometry leads to an increase in the apparent rate of formation of both species (**Figure 5D**). This behavior, which is suggestive of allosteric synergy between substrate and Fe(II) binding, appears to be a common trait of HDOs and supports the assignment of the 625-nm and 510-nm complexes in AetD to μ-peroxodiiron(III) intermediates. Although we have yet to obtain supporting kinetic and spectroscopic evidence, the 510-nm species may represent the second μ-peroxodiiron(III) intermediate that performs the later stage oxidation to yield the nitrile and carbaldehyde products (**Figure 6**).

**Fig. 6.**
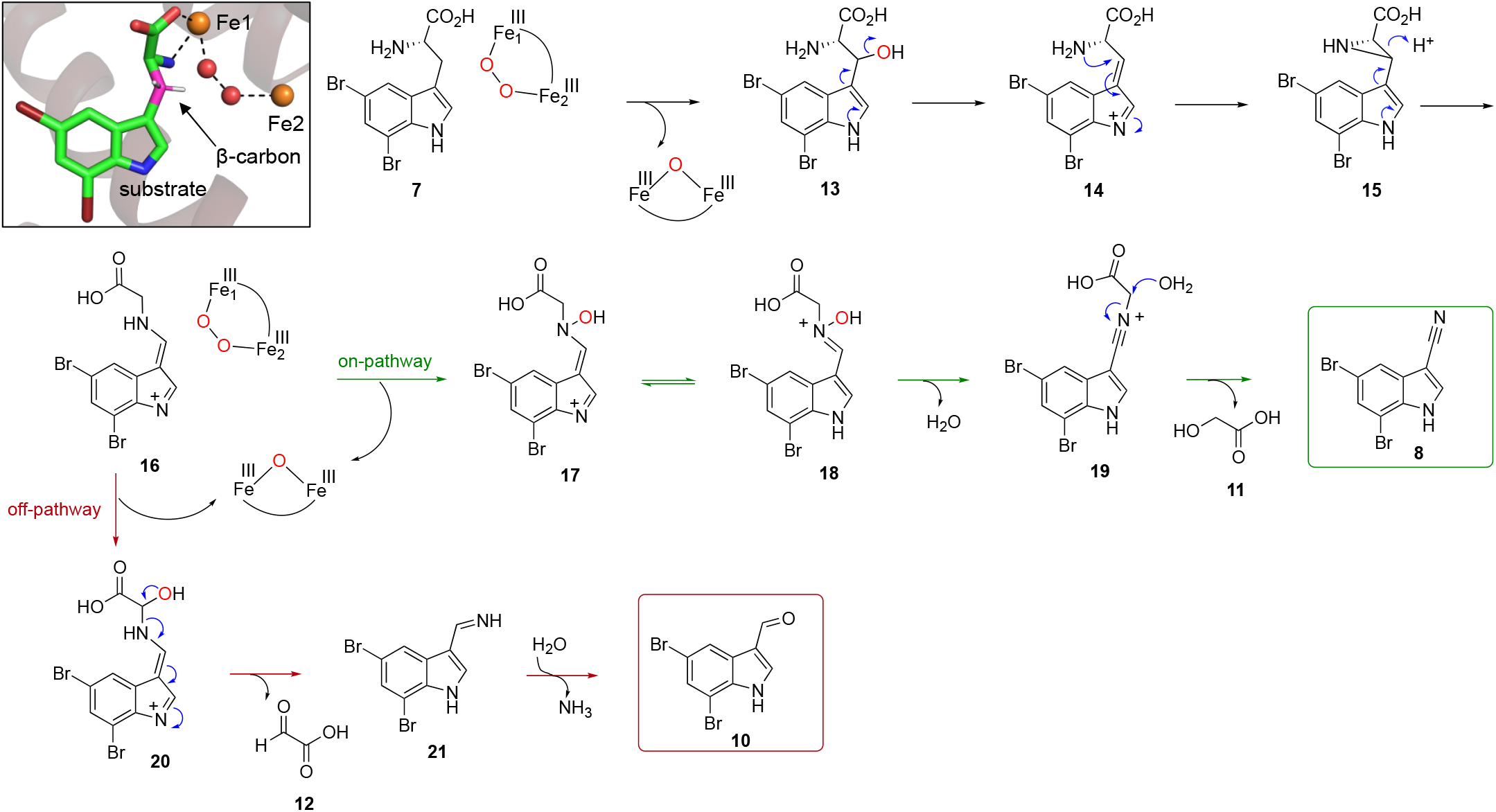
Proposed reaction mechanism of AetD catalyzed reaction. The top left panel is a zoomed-in view of the active site of substrate bound Fe_2_ (II/II)-AetD structure. Substrate β-carbon (colored in pink) is positioned to be hydroxylated by the peroxo-diiron intermediate (peroxo species modeled in as red spheres). Hydrogens on the β-carbon are modeled as white sticks. Orange spheres are Fe ions.

## Discussion and conclusion

AetD represents a new addition to the HDO superfamily and performs a challenging rearrangement reaction yielding a nitrile product, providing an alternative route for nitrile biosynthesis, and expanding the chemistry performed by HDO enzymes. In terms of the mechanism of nitrile formation by AetD, our structural data indicate that substrate binds prior to diiron cofactor formation, as has been proposed for BesC^27,28^ and UndA^26^. In AetD, the substrate-binding pocket is only accessible in the apo state when side chains are not positioned for metal binding and core α3 is unwound. Our snapshots further suggest that substrate-binding repositions key histidine side chains for Fe1 binding, and that Fe1 binding in turn positions glutamate and aspartate side chains for Fe2 binding. Fe1 and Fe2 can access the cofactor-binding sites through unwound turn of core α3, which appears to gate substrate entry and cofactor assembly as previously observed in HDO enzyme SznF.^25^ An excess iron (Fe3) is observed in what appears to be the entry route. It is only when core α3 is most ordered that the diiron cofactor obtains its full complement of ligands with the highly flexible core α3 residue H179 providing the final ligation to complete cofactor assembly.

Our SF-absorption spectra support the prediction from structure that substrate must bind first, showing that formation of the peroxodiiron(III) intermediate is substrate-triggered. This observation is in contrast to SznF, for which spectroscopic data show that substrate binding is independent of dioxygen addition and peroxo-adduct formation.^24^ The latter result for SznF suggests that diiron cofactor assembly can precede substrate binding, which explains why a crystal structure of substrate-free diiron-SznF could be attained^25^ whereas a structure of substrate-free diiron-AetD could not.

Our structural and mechanistic studies allow us to propose that the AetD reaction proceeds via initial hydrogen atom abstraction and hydroxylation at the β-carbon of 5,7-dibromo-L-tryptophan **(Figure 6**). We find that the coordination of the substrate to Fe1 positions the β-carbon right next to an open site on the diiron cofactor that must be the binding site for dioxygen (**Figure 6 inset**). The transient absorption spectra of AetD are rich in features and best described considering formation of two peroxodiiron(III) species that are involved in the two oxidative events to yield the nitrile product. Assignment of these species are consistent with their optical features, kinetics, O_2_- and Fe-dependence, but their precise characterization requires additional studies utilizing Mössbauer spectroscopy, isotopically labeled substrates and proposed intermediates (**16**), which are under way. Assuming for the time being that two peroxodiiron(III) species are involved, we can propose that the 625-nm species carries out the μ-peroxodiiron(III)-mediated hydroxylation at the β-carbon of Trp, which would form compound **13**. Loss of water gives the exocyclic double bond containing compound **14** that in turn is trapped by the amine group to form the aziridine intermediate **15**.

Alternatively, following the example of the Fe(II)/α-ketoglutarate dependent oxygenase TqaL during 2-aminoisobutyrate biosynthesis in fungi, the aziridine could form via carbocation-mediated chemistry (**Figure S14**).^38^ Regardless of the precise event, aziridine ring opening in which the nitrogen atom is inserted between the α and β carbons gives intermediate **16**. We believe that this intermediate is the branching point between on and off-pathway reactions. The on-pathway reaction likely proceeds via a second μ-peroxodiiron(III)-mediated N-hydroxylation (510-nm species) to **17** and tautomerization to yield nitrone intermediate **18**. Formation of the second proposed μ-peroxodiiron(III) is likely afforded by the excess ascorbate present that allows for a subsequent reaction of the regenerated diferrous center with **16** and molecular oxygen. Formation of two peroxo(III/III) adducts is not entirely unprecedented and has been previously reported for the N-oxygenase SznF,^24^ for which, however, these species are spectroscopically indistinguishable and not ‘substrate’-triggered, in contrast to AetD. Loss of water to **19** allows for the re-addition of water at the α-carbon to release glycolic acid (the product as established here) and give the nitrile product **8**. In the shunt off-pathway, intermediate **16** rather undergoes C-hydroxylation to yield intermediate **20**. C–N bond cleavage results in the formation of imine intermediate **21** and glyoxylic acid. Upon spontaneous hydrolysis, the imine gives 5,7-dibromo-indole-3-carbaldehyde **10** as the shunt product.

Our overall mechanistic proposal for an amine nitrogen migration to a β-carbon has literature precedence. Three different classes of aminomutases are known to carry out such reactions^39^: adenosylcobalamin (B_12_)-dependent aminomutases (e.g. lysine 5,6-aminomutase), *S*-adenosyl methionine (AdoMet)-dependent aminomutases (e.g. lysine 2,3-aminomutase), and methylideneimidazole-5-one (MIO)-dependent aminomutases (e.g. phenylalanine aminomutase). In the case of the AetD catalyzed reaction, in addition to amine nitrogen migration to the β-carbon, there is also C_α_-C_β_ bond cleavage and additional oxidation. A single enzyme catalyzed C_α_-C_β_ bond cleavage reaction on tryptophan is rare with the radical *S*-adenosyl methionine-dependent tryptophan lyase NosL being the lone example.^40,41^ NosL catalyzes a similar bond cleavage reaction during nosiheptide antibiotic biosynthesis while converting L-tryptophan to 3-methylindole-2-carboxylic acid. It is quite remarkable that AetD singlehandedly carries out both steps which otherwise would have been a multi-enzyme catalyzed process.

In addition to their chemical prowess, AetD and the other members of the HDO superfamily also exemplify the different ways by which nature tinkers with enzyme architectures (conformational gating) and substrate properties (metal-binding ability) to diversify and control enzymatic activities. Although we are at a nascent stage of discovering and exploring the wealth of the reactivities performed by HDOs, the results to date strongly argue that HDOs may be far more diverse with respect to structure and reactivity when compared to FDOs, which may be in part attributed to the apparent plasticity of their cofactors. We anticipate that this work will inspire future efforts of rational design and directed evolution on AetD and other enzymes in the HDO superfamily, by taking advantage of their cofactors and dynamic scaffolds to evolve new chemical reactivities and/or expanded substrate scope. For example, engineering AetD to allow for incorporation of nitrile functional groups into other amino acid substrates is an exciting direction towards expanding our inventory of nitrile containing compounds.

## Acknowledgments

Funding was generously provided by the National Institutes of Health (R01-ES030316 to B.S.M., F32-ES033540 to A.L.L., R35-GM126982 to C.L.D., and GM111978 and GM126303 to M.-E.P.), the National Science Foundation (OCE-1837116 to B.S.M.), and the Swiss NSF postdoctoral fellowship (P2EZP3_195643 to R.J.B.S). We thank Tadhg P. Begley and Avick Kumar Ghosh (Texas A&M University) for providing us the tryptophan synthase overexpression construct and for insightful discussion regarding the mechanism of AetD catalyzed reaction. C.L.D. is a Howard Hughes Medical Institute Investigator. This work is based upon research conducted at the Northeastern Collaborative Access Team beamlines, which are funded by the National Institute of General Medical Sciences from the National Institutes of Health (P30-GM124165). This research used resources of the Advanced Photon Source, a U.S. Department of Energy (DOE) Office of Science User Facility operated for the DOE Office of Science by Argonne National Laboratory under Contract No. DE-AC02-06CH11357. Use of the Stanford Synchrotron Radiation Lightsource, SLAC National Accelerator Laboratory, is supported by the U.S. DOE, Office of Science, Office of Basic Energy Sciences under Contract No. DE-AC02-76SF00515. The SSRL Structural Molecular Biology Program is supported by the DOE Office of Biological and Environmental Research, and by the National Institutes of Health (P30-GM133894).

## References

1 Fleming, F. F., Yao, L., Ravikumar, P. C., Funk, L. & Shook, B. C. Nitrile-Containing Pharmaceuticals: Efficacious Roles of the Nitrile Pharmacophore. J. Med. Chem. 53, 7902–7917, (2010).

2 Wang, X. et al. Nitrile-containing pharmaceuticals: target, mechanism of action, and their SAR studies. RSC Med. Chem. 12, 1650–1671, (2021).

3 Pollak, P., Romeder, G., Hagedorn, F. & Gelbke, H.-P. in Ullmann’s Encyclopedia of Industrial Chemistry (2000).

4 F. Fleming F. Nitrile-containing natural products. Nat. Prod. Rep. 16, 597–606, (1999).

5 Scheuer, P. J. Isocyanides and cyanides as natural products. Acc. Chem. Res. 25, 433–439, (1992).

6 Legras, J. L., Chuzel, G., Arnaud, A. & Galzy, P. Natural nitriles and their metabolism. World J. Microbiol. Biotechnol. 6, 83–108, (1990).

7 Duffy, S. S. Cyanide in Biology. (Academic Press, London, 1981).

8 Kahn, R. A., Bak, S., Svendsen, I., Halkier, B. A. & Møller, B. L. Isolation and reconstitution of cytochrome P450ox and in vitro reconstitution of the entire biosynthetic pathway of the cyanogenic glucoside dhurrin from sorghum. Plant Physiol. 115, 1661–1670, (1997).

9 Nomura, J. et al. Crystal structure of aldoxime dehydratase and its catalytic mechanism involved in carbon-nitrogen triple-bond synthesis. Proc. Nat. Acad. Sci. U.S.A 110, 2810–2815, (2013).

10 Konishi, K. et al. Discovery of a reaction intermediate of aliphatic aldoxime dehydratase involving heme as an active center. Proc. Nat. Acad. Sci. U.S.A 103, 564–568, (2006).

11 Nelp, M. T. & Bandarian, V. A Single Enzyme Transforms a Carboxylic Acid into a Nitrile through an Amide Intermediate. Angew. Chem. Int. Ed. 54, 10627–10629, (2015).

12 Lane, A. L. et al. Structures and comparative characterization of biosynthetic gene clusters for cyanosporasides, enediyne-derived natural products from marine actinomycetes. J. Am. Chem. Soc. 135, 4171–4174, (2013).

13 Olano, C. et al. Biosynthesis of the angiogenesis inhibitor borrelidin by Streptomyces parvulus Tü4055: insights into nitrile formation. Mol. Microbiol. 52, 1745–1756, (2004).

14 Adak, S., Lukowski, A. L., Schäfer, R. J. B. & Moore, B. S. From Tryptophan to Toxin: Nature’s Convergent Biosynthetic Strategy to Aetokthonotoxin. J. Am. Chem. Soc. 144, 2861–2866, (2022).

15 Breinlinger, S. et al. Hunting the eagle killer: A cyanobacterial neurotoxin causes vacuolar myelinopathy. Science (New York, N.Y.) 371, eaax9050, (2021).

16 Hull, A. K., Vij, R. & Celenza, J. L. Arabidopsis cytochrome P450s that catalyze the first step of tryptophan-dependent indole-3-acetic acid biosynthesis. Proc. Nat. Acad. Sci. U.S.A 97, 2379–2384, (2000).

17 Rajakovich, L. J. et al. in Comprehensive Natural Products III (eds Hung-Wen Liu & Tadhg P. Begley) 215–250 (Elsevier, 2020).

18 Ng, T. L., Rohac, R., Mitchell, A. J., Boal, A. K. & Balskus, E. P. An N-nitrosating metalloenzyme constructs the pharmacophore of streptozotocin. Nature 566, 94–99, (2019).

19 Hedges, J. B. & Ryan, K. S. In vitro Reconstitution of the Biosynthetic Pathway to the Nitroimidazole Antibiotic Azomycin. Angew. Chem. Int. Ed. 58, 11647–11651, (2019).

20 Patteson, J. B. et al. Biosynthesis of fluopsin C, a copper-containing antibiotic from Pseudomonas aeruginosa. Science (New York, N.Y.) 374, 1005–1009, (2021).

21 Rui, Z. et al. Microbial biosynthesis of medium-chain 1-alkenes by a nonheme iron oxidase. Proc. Nat. Acad. Sci. U.S.A 111, 18237–18242, (2014).

22 Marchand, J. A. et al. Discovery of a pathway for terminal-alkyne amino acid biosynthesis. Nature 567, 420–424, (2019).

23 Manley, O. M. et al. Self-sacrificial tyrosine cleavage by an Fe:Mn oxygenase for the biosynthesis of para-aminobenzoate in Chlamydia trachomatis. Proc. Nat. Acad. Sci. U.S.A 119, e2210908119, (2022).

24 McBride, M. J. et al. A Peroxodiiron(III/III) Intermediate Mediating Both N-Hydroxylation Steps in Biosynthesis of the N-Nitrosourea Pharmacophore of Streptozotocin by the Multi-domain Metalloenzyme SznF. J. Am. Chem. Soc. 142, 11818–11828, (2020).

25 McBride, M. J. et al. Structure and assembly of the diiron cofactor in the heme-oxygenase– like domain of the N-nitrosourea–producing enzyme SznF. Proc. Nat. Acad. Sci. U.S.A 118, e2015931118, (2021).

26 Zhang, B. et al. Substrate-Triggered Formation of a Peroxo-Fe2(III/III) Intermediate during Fatty Acid Decarboxylation by UndA. J. Am. Chem. Soc. 141, 14510–14514, (2019).

27 Manley, O. M. et al. BesC Initiates C–C Cleavage through a Substrate-Triggered and Reactive Diferric-Peroxo Intermediate. J. Am. Chem. Soc. 143, 21416–21424, (2021).

28 McBride, M. J. et al. Substrate-Triggered μ-Peroxodiiron(III) Intermediate in the 4-Chloro-l-Lysine-Fragmenting Heme-Oxygenase-like Diiron Oxidase (HDO) BesC: Substrate Dissociation from, and C4 Targeting by, the Intermediate. Biochemistry 61, 689–702, (2022).

29 Schwarzenbacher, R. et al. Structure of the Chlamydia Protein CADD Reveals a Redox Enzyme That Modulates Host Cell Apoptosis. J. Biol. Chem. 279, 29320–29324, (2004).

30 Andrews, S. C. The Ferritin-like superfamily: Evolution of the biological iron storeman from a rubrerythrin-like ancestor. Biochim. Biophys. Acta 1800, 691–705, (2010).

31 Manley, O. M., Fan, R., Guo, Y. & Makris, T. M. Oxidative Decarboxylase UndA Utilizes a Dinuclear Iron Cofactor. J. Am. Chem. Soc. 141, 8684–8688, (2019).

32 Kawasaki, H., Bauerle, R., Zon, G., Ahmed, S. A. & Miles, E. W. Site-specific mutagenesis of the alpha subunit of tryptophan synthase from Salmonella typhimurium. Changing arginine 179 to leucine alters the reciprocal transmission of substrate-induced conformational changes between the alpha and beta 2 subunits. J. Biol. Chem. 262, 10678–10683 (1987).

33 Böttcher, C. et al. The Biosynthetic Pathway of Indole-3-Carbaldehyde and Indole-3-Carboxylic Acid Derivatives in Arabidopsis. Plant Physiol. 165, 841–853, (2014).

34 Zelante, T. et al. Tryptophan catabolites from microbiota engage aryl hydrocarbon receptor and balance mucosal reactivity via interleukin-22. Immunity 39, 372–385, (2013).

35 Roager, H. M. & Licht, T. R. Microbial tryptophan catabolites in health and disease. Nat. Commun. 9, 3294, (2018).

36 Olguin-Uribe, G. et al. 6-Bromoindole-3-Carbaldehyde, from an Acinetobacter Sp. Bacterium Associated with the Ascidian Stomozoa murrayi. J. Chem. Ecol. 23, 2507–2521, (1997).

37 Park, K. et al. Peroxide Activation for Electrophilic Reactivity by the Binuclear Non-heme Iron Enzyme AurF. J. Am. Chem. Soc. 139, 7062–7070, (2017).

38 Cha, L. et al. Mechanistic Studies of Aziridine Formation Catalyzed by Mononuclear Non-Heme Iron Enzymes. J. Am. Chem. Soc. 145, 6240–6246, (2023).

39 Wu, B., Szymański, W., Heberling, M. M., Feringa, B. L. & Janssen, D. B. Aminomutases: mechanistic diversity, biotechnological applications and future perspectives. Trends Biotechnol. 29, 352–362, (2011).

40 Sicoli, G. et al. Fine-tuning of a radical-based reaction by radical S-adenosyl-L-methionine tryptophan lyase. Science (New York, N.Y.) 351, 1320–1323, (2016).

41 Bhandari, D. M., Fedoseyenko, D. & Begley, T. P. Mechanistic Studies on Tryptophan Lyase (NosL): Identification of Cyanide as a Reaction Product. J. Am. Chem. Soc. 140, 542–545, (2018).

