## SUPPLEMENTARY INFORMATION for "Oxidative rearrangement of tryptophan to indole nitrile by a single diiron enzyme"

### General Considerations

All chemicals were purchased from Sigma-Aldrich unless mentioned otherwise. LB broth (Lennox), TB broth, and kanamycin were purchased from Fisher Scientific. Ampicillin was from EMD Millipore. Potassium phosphate salt was purchased from Teknova. Amicon® Ultra-15 Centrifugal Filter Unit (10,000 MWCO) was obtained from Millipore. HisTrap column was obtained from GE Healthcare. Econo-Pack 10DG desalting columns were purchased from Bio-Rad. HPLC and LC-MS solvents were purchased from Fisher Chemical and were used without further purification. pET-28a(+) plasmid containing *E. coli* codon-optimized *aetD* gene between NdeI and XhoI restriction sites was synthesized and subcloned by Twist Bioscience. NMR spectroscopic data were obtained on a 500 MHz JEOL NMR spectrometer with a 3.0 mm probe. The values of the chemical shifts are described in ppm and coupling constants are reported in Hz. NMR data analysis was performed using MestreNova® 14.21-27684, 2021.

### Preparatory HPLC parameters

Preparatory HPLC was performed with an Agilent Technologies system composed of a PrepStar pump, a ProStar 410 autosampler, a ProStar UV detector, and a fraction collector. A Phenomenex Luna C18 column (5μm, C18 100 Å, 100 x 21.2 mm) was used for separation. Data was collected and analyzed using Agilent OpenLAB CDS ChemStation software version C.01.05.

LC conditions:
A- water + 0.1% formic acid
B- acetonitrile + 0.1% formic acid

HPLC method: 10 mL/min flow rate; 0-5 min 5% B, 5-20 min 100% B, 20-25 min 100%B, 25-28 min 5% B, 28-30 min 5% B

### LC-MS parameters

LC-MS analysis was performed with an Agilent Technologies 1260 Infinity series HPLC equipped with a degasser, binary pump, autosampler, and diode array detector coupled to an Agilent Technologies 6530 Accurate-Mass Q-TOF LC/MS. Separations were achieved using a Kinetex 5 μm C18 100 Å, 150 x 4.6 mm column. Data was collected and analyzed using MassHunter Workstation Software version B.05.01.

LC conditions:
A- water + 0.1% formic acid
B- acetonitrile + 0.1% formic acid

LC method: 0.75 mL/min flow rate; 0-4 min 10% B, 4-11 min 10-95% B, 11-21 min 95% B, 21-23 min 95%-10% B, 23-27 min 10% B. Dual ESI ion source, negative ionization mode.

**Overexpression and purification of AetD**

A pET-28a(+) plasmid containing the *E. coli* codon-optimized *aetD* gene between NdeI and XhoI restriction sites was synthesized and subcloned by Twist Bioscience. For activity assay, AetD overexpression was performed in TB media following the reported protocol.[1] For crystallographic and biophysical experiments, AetD overexpression was carried out in M9 minimal media supplemented with 125 μM (final concentration) of $Fe(NH_4)_2(SO_4)_2$. In both cases, Ni affinity chromatography was used for the purification of the N-terminal His-tagged protein (sequence: MGSSHHHHHHSSGLVPRGSHM) following the reported protocol.[1] An additional round of purification was performed using size exclusion chromatography (Sephadex S200) on a fast protein liquid chromatography (FPLC) conducted on a Cytiva ÄKTA Pure 25 L1 system fitted with an F9-C fraction collector and S9 sample pump and controlled by Unicorn v7 software. AetD was found to be dimer in solution based on the comparison of elution column volume with standards.

**Overexpression and purification of Se-Met AetD**

A 125 mL sterilized flask with 50 mL sterile LB was prepared. Kanamycin was added to the media with a final 40 μg/mL concentration. A single colony of *E. coli* BL21(DE3) harboring the AetD encoding gene in a pET-28a(+) vector was transferred to the media. The cell culture was grown at 37 ºC with shaking at 220 rpm for 16-20 h. The cell culture was pelleted by centrifugation at 4000 x g for 10 min at RT. The cell pellet was resuspended in 20 mL M9-kanamycin media. The cells were pelleted by spinning down. The above step was repeated to remove residual LB. The pellet was resuspended in 1L M9-kanamycin media. Cells were grown at 37 ºC with shaking until the $OD_{600}$ ~ 0.3. L-Lysine, L-Phenylalanine, and L-Threonine were added to the culture with 100 mg/L final concentration while L-Isoleucine, L-Leucine, L-Valine, and L-Selenomethionine were added to the culture with 50 mg/L final concentration. The culture was allowed to reach $OD_{600}$ ~ 0.6 when 0.5 mM IPTG was added. The induced culture was incubated at 15 ºC (220 rpm) for 20 h. Cells were harvested and protein was purified using Ni affinity chromatography following the reported protocol.[1] An additional round of purification was performed using size exclusion chromatography (Sephadex S200) on a fast protein liquid chromatography (FPLC) conducted on a Cytiva ÄKTA Pure 25 L1 system fitted with an F9-C fraction collector and S9 sample pump and controlled by Unicorn v7 software.

**Overexpression of tryptophan synthase**

A stock of *E. coli* BL21 (DE3) containing the overexpression plasmid of tryptophan synthase (pSTB7) was grown overnight in 10 ml LB medium supplemented with 100 µg/ml of ampicillin. 10 ml of this overnight culture was used to inoculate 1.0 L of LB medium containing 100 µg/ml of ampicillin. The cells were grown at 37 °C at 200 rpm for 24 h. The cells were harvested by centrifuging at 9000 rpm for 15 min. The cells were suspended in 35 ml of lysis buffer (100 mM $KH_2PO_4$, pH 7.5). 5 mg of pyridoxal 5'-phosphate (PLP) was added, and 4 cycles of sonication were performed (each cycle is for 30 sec with 1sec on/1sec off, 65% power) at 5 min intervals. The lysate was centrifuged at 18,000 rpm for 20 min at 4 °C and filtered through 0.2 µm filters to remove cell debris. The lysate was stored at 4 °C for up to one month and was used whenever required in this period.

**Enzymatic synthesis of 5,7-dibromo-L-tryptophan**

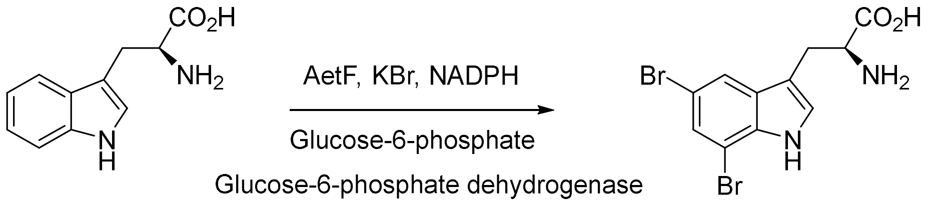

A 10 mL AetF reaction was prepared with 10 µM AetF, 1 U/mL glucose-6-phosphate dehydrogenase, 10 mM glucose-6-phosphate, 1 mM NADPH, 5 mM L-tryptophan (10 mg total), 50 mM KBr, in 100 mM $KP_i$ pH 7.5 buffer. The reaction was incubated at 30 °C, 200 rpm overnight. The reaction mixture was lyophilized overnight and extracted 3 times with 1 mL methanol. The extraction was filtered through an 0.2 µm syringe filter and concentrated to ~500 µL by rotary evaporation. The 5,7-dibromo-L-tryptophan product was purified by preparative HPLC following the protocol described above. The typical yield was 7-8 mg.

**Enzymatic synthesis of $^{13}C$-labeled isotopologues of L-tryptophan**

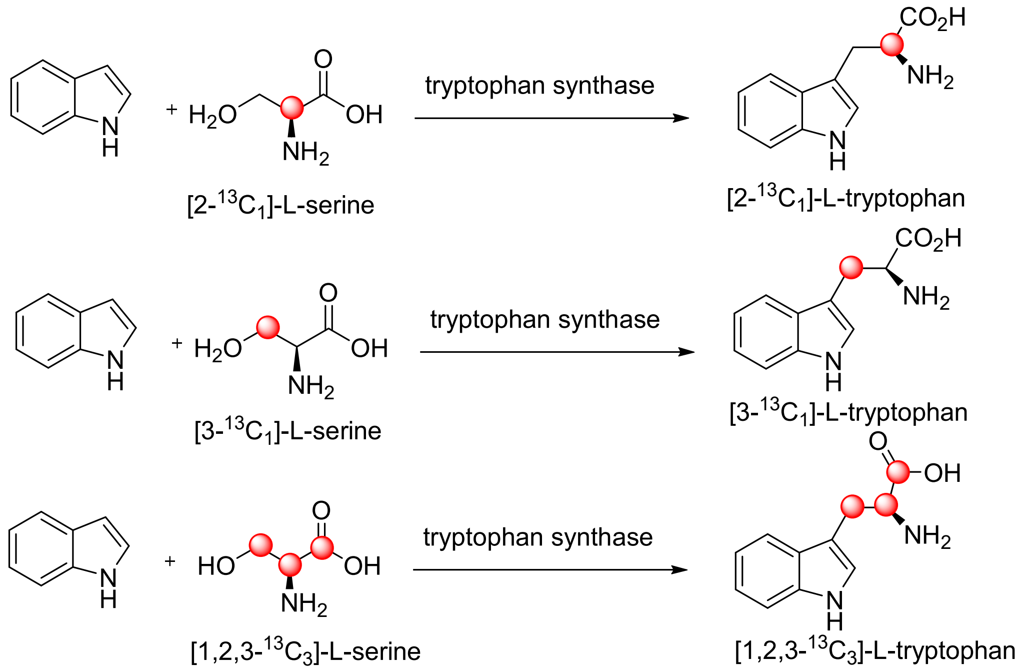

$^{13}C$-Labeled isotopologues of L-tryptophan were produced following the reported protocol[2] with slight modification. 10 mg of indole and 12 mg of the respective $^{13}C$-labeled isotopologues of L-serine were dissolved in 10 ml of 100 mM potassium phosphate buffer (pH 7.5). 1 mL of cell lysate was added to initiate the enzymatic reaction. The reaction mixture was incubated at 37 °C at 200 rpm for 3 h. After the completion of the reaction, the reaction mixture was lyophilized, and the product was redissolved in methanol. Proteins were removed by ultrafiltration using 10 kDa cut-off filters (American Chromatography Supplies). Enzymatic products were purified on preparative HPLC using the protocol described above. The typical yield was 5-7 mg.

**AetD reaction with $^{13}C$ labeled isotopologues of 5,7-dibromo-L-tryptophan**

100 µL reaction consisting of 20 µM AetF, 2 mM $^{13}C$-labeled L-tryptophan, 5 mM NADPH, 20 mM KBr, 1 U/mL glucose-6-phosphate dehydrogenase, 5 mM glucose-6-phosphate in 100 mM $KP_i$ pH 7.5 buffer was prepared in 1.5 mL centrifuge tubes and incubated at RT for 6 h. Then AetD,

Fe (II), and ascorbate were added to the reaction mixture with final concentrations of 250 µM, 5mM, and 5mM respectively and the final volume of the reaction mixture was 200 µL. The resulting reaction mixture was incubated at RT overnight. The samples were analyzed using LC-MS and $^{13}$C-NMR spectroscopy. For $^{13}$C-NMR experiments, the samples were transferred to a 3mm NMR tube after adding 10% $D_2O$.

**Derivatization of glyoxylic acid formed in AetD reaction using o-phenylenediamine**

A 100 µL reaction mixture containing 500 µM 5,7-dibromo-L-tryptophan, 40 µM AetD, and 5mM Fe (II) was incubated at RT overnight. Next, the assay solution was made acidic (pH=1) by adding HCl. Then, o-phenylenediamine was added to a final concentration of 1 mM. The assay mixture was kept at room temperature for 2h. The reaction was quenched by adding 1:1 methanol, and the resulting solution was filtered using 0.2 µM centrifugal filters. The samples were analyzed by LC-MS.

**Stopped flow protocol**

Stopped-flow Absorption (SF-Abs) measurements were carried out in a SX20 stopped-flow spectrophotometer from Applied Photophysics Ltd. (Leatherhead, UK) that is housed in an anoxic chamber (Coy Laboratories, Michigan, MI). All experiments were carried out 5 °C and were single-mix (two-syringe) experiments, in which the AetD reactant solution was diluted by 2-fold. An $O_2$-free solution of AetD (0.30 mM) that was reconstituted with two molar equivalents of Fe(II) (0.6 mM), with or without the dibromo-Tryptophan substrate, was mixed with $O_2$-saturated (on ice) 50 mM sodium HEPES buffer (pH 7.5) containing 10% glycerol, giving an estimated $O_2$ concentration after mixing of 0.9 mM. All time-resolved absorption spectra were recorded with the photodiode array (PDA) detector. In experiments in which we examined the dependence of the formation of the intermediate(s) as a function of Fe equivalents, the $O_2$-free AetD was mixed with different Fe amounts prior to reaction with the $O_2$-saturated buffer. The kinetics of formation and decay of the intermediates were fit by linear regression using equation 1:

$$\Delta A_{int}(t) = \frac{A\,k_1}{k_2 - k_1}\left(e^{-k_1 t} - e^{-k_2 t}\right)$$

Because the kinetic traces suggest that the intermediates exist in two forms, in most cases for fitting of the data we used the more expanded form to include an additional term accounting for the fact that the decay of the intermediates to the following state occurs with two different rate constants:

$$\Delta A_{int}(t) = \frac{A\,k_1}{k_2 - k_1}\left(e^{-k_1 t} - e^{-k_2 t}\right) + \frac{A'\,k_1}{k_2' - k_1}\left(e^{-k_1 t} - e^{-k_2' t}\right)$$

**Crystallization**

All crystallization experiments were performed in an MBraun anaerobic chamber in an $N_2$ environment. To prepare substrate-bound protein samples, 5 molar equivalents (1.75 mM) of enzymatically synthesized substrate 5,7 dibromo tryptophan (see page 5) dissolved in DMSO was added to 10.6 mg/mL (0.35 mM, measured by absorbance at 280 nm using UV-Vis spectrophotometer using extinction coefficient of 35410 $M^{-1}$ $cm^{-1}$ calculated by ProtParam Tool, https://web.expasy.org/protparam) AetD or 10.0 mg/mL (0.33 mM) Se-Met labeled AetD in storage buffer (50 mM Tris-HCl, 100 mM NaCl, 10% v/v glycerol, pH 8.0), and incubated overnight at 4 °C. The AetD protein used had an uncleaved N-terminal purification tag with sequence: MGSSHHHHHHSSGLVPRGSHM. Initial crystallization conditions for substrate co-crystallized AetD were identified using crystallization screens dispensed by Mosquito liquid-handling robot (SPT Labtech) at room temperature. The identified crystallization conditions were further optimized using hanging-drop vapor diffusion method in the same anerobic, room-temperature settings. The substrate-bound structure was obtained from AetD expressed and purified from M9 minimal media without iron supplementation. The structure with metallocofactor partially-reconstituted was obtained from AetD expressed and purified from M9 minimal media supplemented with 125 μM (final concentration) of $Fe(NH_4)_2(SO_4)_2$. Crystals that yielded these two structures grew in the following crystallization condition: 100 mM 2-(N-Morpholino) ethanesulfonic acid (MES) monohydrate at pH 6.0, 20% w/v polyethylene glycol (PEG) 4,000, and a salt mixture at pH 6.0 (110 mM malonic acid, 15 mM ammonium citrate tribasic, 7.2 mM succinic acid, 18 mM DL-malic acid, 24 mM sodium acetate trihydrate, 30 mM sodium formate, 9.6 mM ammonium tartrate dibasic). The structure with the fully-reconstituted metallocofactor was obtained from AetD expressed and purified from M9 minimal media without iron supplementation, and crystal grew in the following crystallization condition: 250 mM ammonium sulfate and 20% w/v PEG 3,350. These crystals were then soaked in 20 mM (final concentration) Fe(II) by adding to the drops an equal volume of well solution supplemented with 40 mM $Fe(NH_4)_2(SO_4)_2$ and 10 mM substrate followed by gentle mixing and overnight incubation. All chemicals used for crystallization were purchased from Hampton Research. To reconstitute Fe(II) at the enzyme active site, selected crystals were soaked in Fe(II) by adding to the drops an equal volume of well solution supplemented with 40 mM $Fe(NH_4)_2(SO_4)_2$ and 10 mM substrate followed by gentle mixing and overnight incubation. The hanging drops consisted of 1.0 µL of AetD or Se-Met labeled AetD co-crystallized with substrate, and 2.0 µL of well solution from one of the two conditions above, in a sealed well over 500 µL of well solution. Transparent plate, rod, or hexagonal-prism shaped crystals typically appeared overnight and grew to full size within 7 days. Crystals used for structure determinations were transferred to a Coy anaerobic chamber with an Ar/$N_2$ gas mix environment for harvesting. Crystals were harvested, cryoprotected with either paraffin oil or the crystallization well solution supplemented with 20% v/v glycerol, and flash frozen in liquid nitrogen.

**X-ray data collection and processing**

X-ray diffraction data were collected at cryogenic temperature (100K) at the Advanced Photon Source (Argonne, IL) beamline 24-ID-C of the Northeastern Collaborative Access Team (NE-CAT) using an Dectris Eiger-2 X 16 M detector, and at the Stanford Synchrotron Radiation Lightsource (Menlo Park, CA) beamline 9–2 using a Dectris Pilatus 6 M detector. The inverse-beam method was used to collect Fe and Se peak data in six 60$^{\circ}$ wedges to capture anomalous signals. All data were indexed, integrated, and scaled in XDS,[3] with data collected at the Fe and Se peak wavelengths processed anomalously with Friedel pairs kept separate. Resolution cutoffs were determined based on considerations of $R_{sym}$, $CC_{1/2}$, I/σ, and data completeness in the highest resolution bin. Data collection statistics are summarized in Table S1.

**Structure determination and refinement**

The initial phases were obtained experimentally from a Se-Met AetD crystal (grown in the first crystallization condition) via single wavelength anomalous dispersion (SAD) method using data collected at Se peak (0.9791 Å). The Se anomalous signal extends to 2.6 Å resolution, where it has a $CC_{1/2\text{-anom}}$ of 0.5 as calculated by Phenix Xtriage.[4] Phenix Hybrid Substructure Search (HySS)[5,6] was used to find the positions of 10 Se sites out of 10 predicted sites for the two molecules in the asymmetric unit (ASU) (excluding the two Met residues that are part of the N-terminus purification tag, see sequence above). Refinement of heavy-atom sites and SAD phasing were performed by Phenix Experimental Phasing (Phaser-EP).[7] An initial experimental map was obtained to 2.8 Å resolution by Phaser-EP with a figure of merit (FOM) of 0.467 before any density modification. This initial map was then subjected to density modifications by Phenix

RESOLVE[8] for solvent flattening and two-fold non-crystallographic symmetry (NCS) averaging, with phases extended to 2.3 Å resolution. An AlphaFold2[9]-generated AetD model was used to generate the mask for NCS averaging. The density-modified experimental map was easily interpretable, providing clear and contiguous electron density for manual model building in Coot.[10] This initial unrefined model in space group $P2_12_12_1$ was used to solve the two other structures of AetD in the same $P2_12_12_1$ space group, i.e. the structures of of AetD with a fully or partially reconstituted cofactor, starting with rigid-body refinements in phenix.refine. 5% of total reflections from the AetD cofactor partially reconstituted dataset were randomly selected and set aside for the test set ($R_{free}$). This test set was carried over for the refinement of the Fe(II) fully-reconstituted structure. The substrate-bound-only structure was solved by molecular replacement (MR) in phenix.phaser using one chain of the initial unrefined model of the Se-Met dataset as the search model without further modifications. A new set of $R_{free}$ was generated for this dataset using 5% randomly selected reflections. Iterative rounds of positional and individual B-factor refinements were performed in phenix.refine, with manual adjustments in Coot against $2|F_o|–|F_c|$ maps contoured to 1.0σ. For the structures solved in space group $P2_12_12_1$ with two molecules in the ASU (see Table S1), two-fold NCS restraints were used throughout the refinements. Structure and geometric restraints for the substrate molecule were generated using eLBOW.[11] $2mF_o$-$DF_c$ composite omit maps were generated, with 5% of the model omitted for individual omit maps, using Phenix Composite_omit_map[12] after each round of refinement to verify the refined structure. Water and other solvent molecules from crystallization conditions were added manually throughout model building using $|F_o|–|F_c|$ map contoured to 3.0σ and the composite omit map contoured to 1.0σ as criteria. Final rounds of refinement included translation, libration, screw (TLS) parameterization with one TLS group per chain in the ASU.

For the substrate-bound structure, the one protomer in the ASU contains all residues in the sequence, 1–239 (out of 239) and 7 residues of N-terminal purification tag. Residues 176 – 183 were missing from the disordered region of helix α3, where no density was observed. The model additionally contained 99 water molecules and one D-malate ion.

For the substrate-bound structure with a partially-reconstituted cofactor, Chain A contains residues 1–238 (out of 239) and 1 residue of the N-terminal purification tag, and chain B contained residues 1–177 and 184–238 (out of 239) and two residues of the N-terminal purification tag. The missing residues (178–183) were from the disordered flexible region of helix α3. The structural model also contained per ASU 169 water molecules, one succinate ion, two D-malate ions, and one glycerol molecule. Following initial rounds of refinements, unaccounted for electron density in $2|F_o|–|F_c|$ maps calculated from native datasets was shown near the active site metal coordination spheres. Anomalous difference maps calculated from datasets collected at the Fe peak wavelength (1.7340 Å) showed consistent high σ electron density peaks: one at Fe in site 1 of chain A (~13 I/α), one at Fe in site 1 of chain B (~12.5 I/α), and one weaker peak at Fe in site 2 of chain B (~ 4.2 I/α), indicating only partial occupancy. Fe ions were modeled in Coot into these electron density peaks. The structure contained three Fe ions (one in the active site of chain A and two in chain B). In chain B, the active site harbors two Fe ions for which the refined crystallographic B-factors are higher than that of Fe1 in chain A when all three Fe ions are set to full occupancy, suggesting partial occupancy of the two iron ions in chain B. To estimate iron occupancy, we set the B-factors of both iron in chain B to be the same as Fe1 in chain A, thereby calculating relative iron occupancy when Fe1 in chain A is assumed to be fully occupied. Refined occupancy shows that Fe1 in chain B and chain A have a similar high occupancy (~84%) and have the same ligands and coordination geometry. Fe2 has almost half of the occupancy (~43%). The identity of two irons in this chain was also confirmed by iron anomalous difference maps (see Figure 3).

For the substrate-bound structure with the fully-reconstituted cofactor, Chain A contains residues 1–238 (out of 239) and six residues of the N-terminal purification tag, and chain B contains residues 1–179 and 185–238 (out of 239) and three residues of the N-terminal purification tag. The missing residues (180–184) were from the disordered flexible region of helix α3 where no density could be observed. The model additionally contained 261 water molecules. Unaccounted for, high σ electron density in $2|F_o|–|F_c|$ maps calculated from native datasets was observed in the cofactor site. Fe anomalous difference map shows consistent high-intensity peaks. Three Fe(II) ions were modeled in chain A and two were modeled in chain B. Iron occupancy was also estimated using the approach described above, where Fe1 in chain A is assumed to be full occupancy.

A summary of the data collection and refinement statistics for all reported structures can be found in Table S1. All structural figures were generated using the PyMOL Molecular Graphics software package (Schrödinger, LLC). Crystallography and AlphaFold packages were compiled by SBGrid.[13]

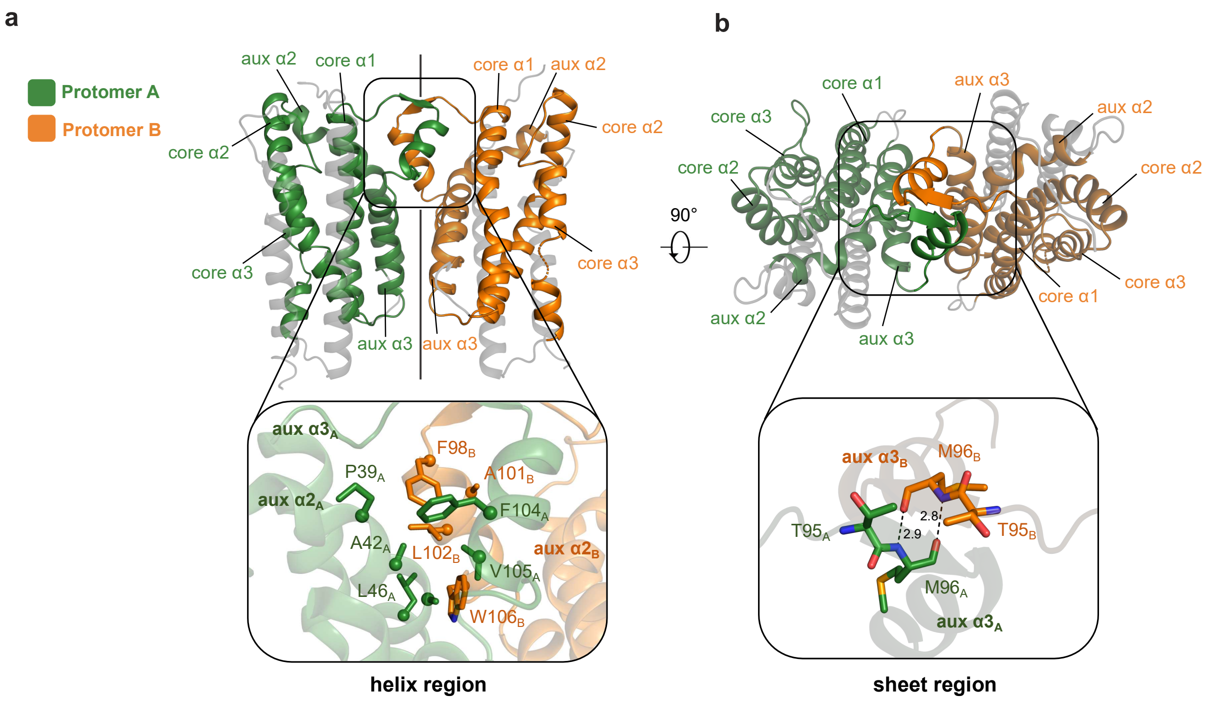

**Fig. S1 Dimeric structure of AetD and the dimer interface. a,** (top) The dimeric structure of AetD. Part of the auxiliary helix α3 forms the dimerization domain (enclosed by the black the box) that consists of a helix region and a sheet region. (bottom) zoomed-in view of the helix region. Only half of this 2-fold symmetrical dimer interface is shown for simplicity. The helix region is largely hydrophobic, containing hydrophobic residues from the dimerization domain and auxiliary helix α2 from both protomers. **b,** the dimeric structure rotated 90° out of plane highlighting the sheet region of the dimerization domain. (bottom) zoomed-in view of the sheet region, which is made of two residues from each protomer (T95 and M96) held together by backbone hydrogen bonds.

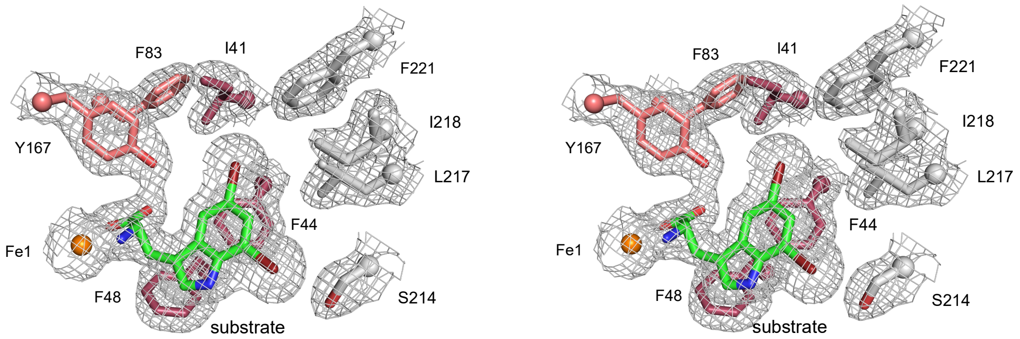

**Fig. S2 Stereoview of substrate electron density**. $2mF_o$-$DF_c$ composite omit maps are contoured at 1.0σ. The final refined model is superimposed. The image on the right is rotated 6 degrees from the left image.

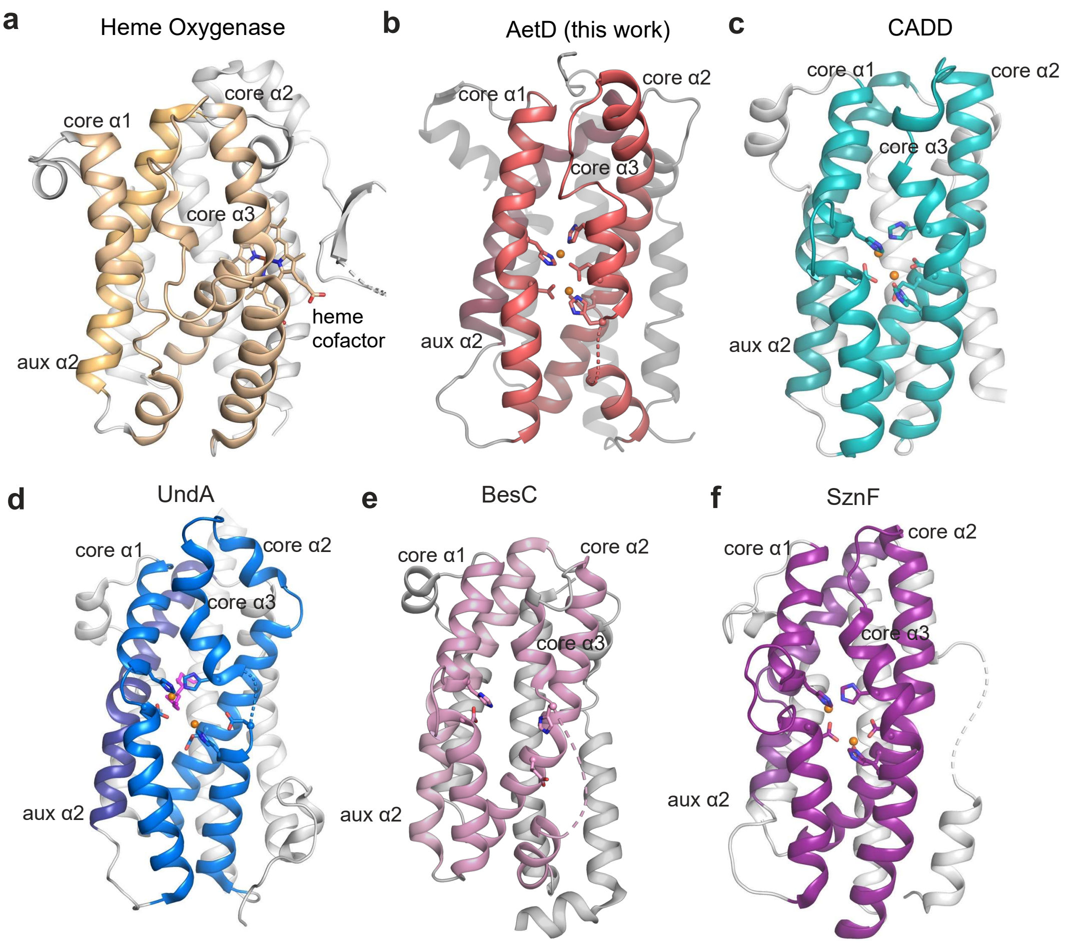

**Fig. S3 Structural comparison of AetD with Heme Oxygenase and other HDO enzymes.** All HDO enzymes share the common architecture of three core helices containing metal-binding ligands, and four auxiliary helices. **a,** Heme Oxygenase[14] (PDB ID: 1WOW) **b,** AetD structure presented in this work. **c,** CADD[15] (PDB ID: 1RCW). **d,** UndA[16] (PDB ID: 6P5Q). **e,** BesC[17] (PDB ID: 7TWA). **f,** SznF[18] (PDB ID: 6VZY).

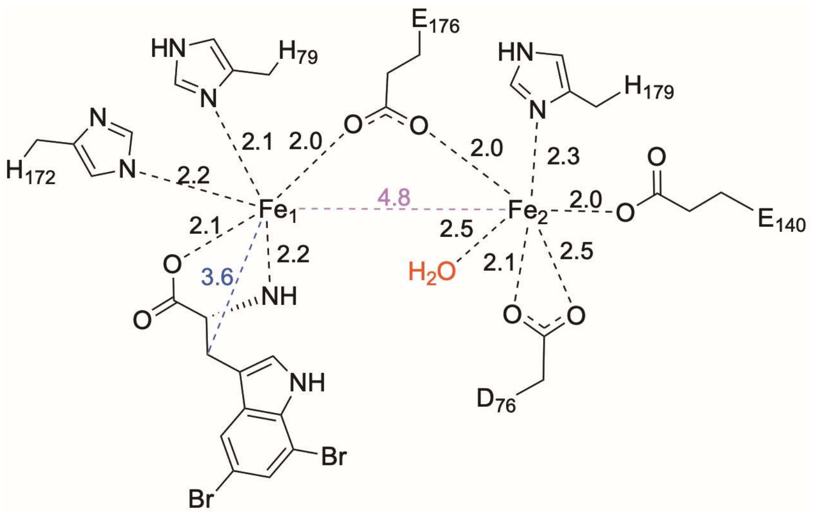

**Fig. S4 Schematic representation of the first coordination sphere of the diiron active site of $Fe_2$(II/II)-AetD.** Fe-ligand distances are in black dashes, Fe-Fe distance is in purple dash, and Fe-substrate (β-carbon) distance is shown in blue dash. All distances are in angstroms (Å).

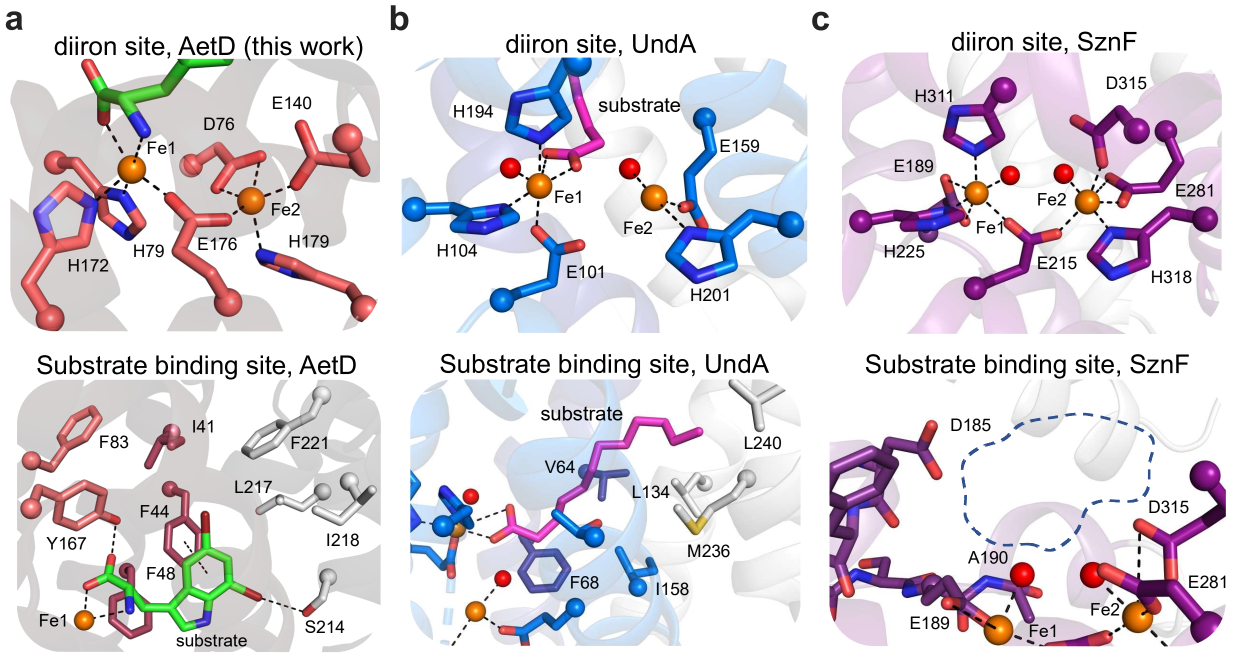

**Fig. S5 HDO structural comparison of substrate binding site and diiron cofactor sites. a,** Diiron site and substrate-binding sites of AetD reproduced here from Figure 3A. Substrate shown with green carbons. **b**, Diiron site and substrate-binding sites of UndA[16] (PDB ID: 6P5Q) are similar to those of AetD. In both structures, the substrate is stabilized by hydrophobic interactions at enzyme core. AetD substrate binds Fe1 in a bidentate manner whereas UndA binds Fe1 only through one of the oxygen atoms in its carboxylate moiety. **c,** Diiron site and putative substrate-binding site of HDO enzyme SznF[18] (PDB ID: 6VZY). Note: no substrate-bound structure is available for SznF; the dashed line in the bottom panel indicates the expected substrate binding position.

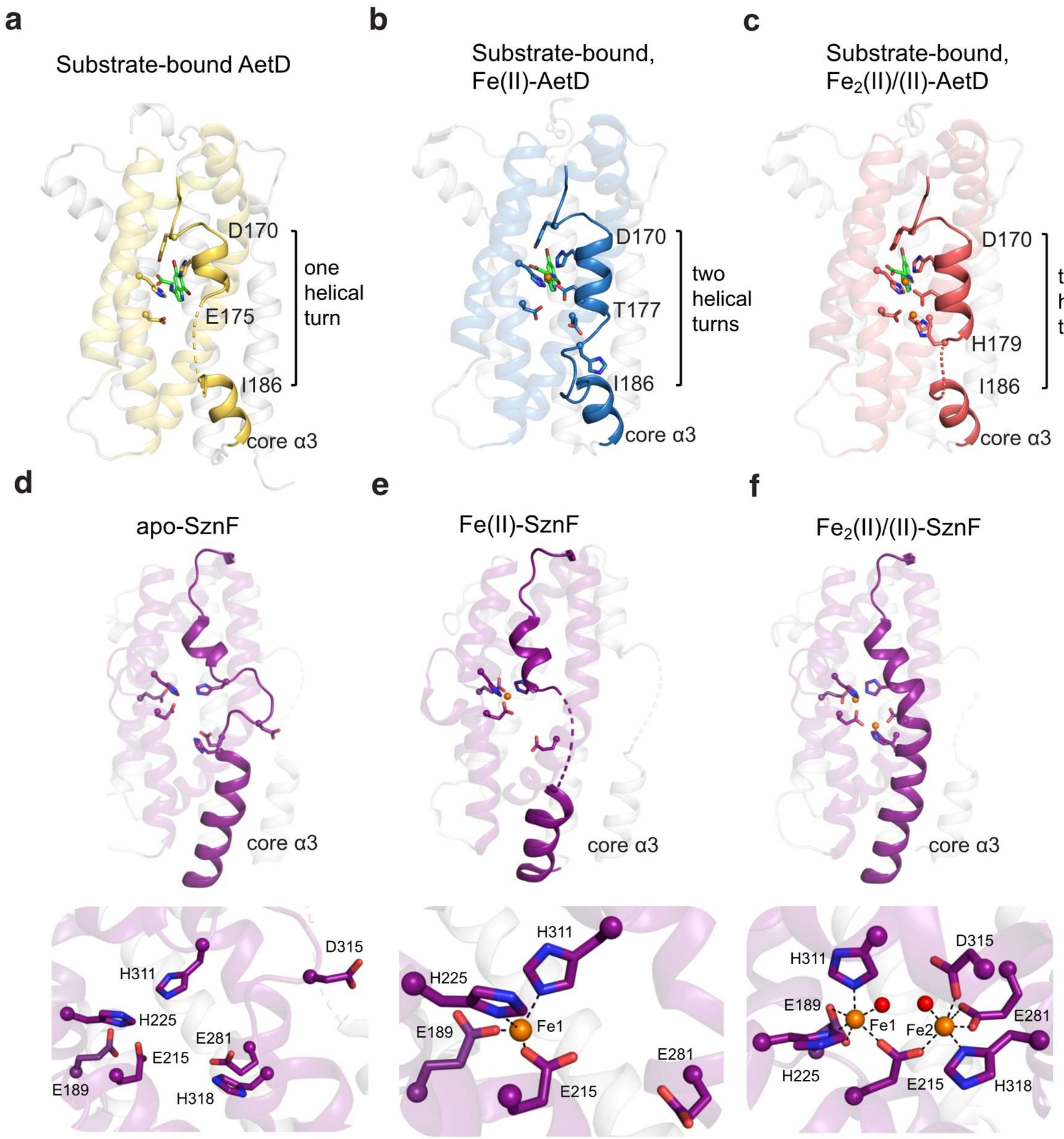

**Fig. S6 Diiron cofactor assembly is conformationally gated by core helix α3. a-c,** Core helix α3 becomes more ordered as the diiron cofactor is assembled. Segment containing residues D170 to H179 presents more ordered secondary structure as shown in increasing number of helical turns. **d-f,** Ordering of core helix α3 is also observed in substrate-independent SznF. When no iron or only one iron is bound at the cofactor, the helix is partially disordered; when both irons are bound, the entire helix becomes fully ordered. PDB IDs of different SznF structures are: **d,** 6M9R[19], **e,** 6M9S[19], and **f,** 6VZY.[18]

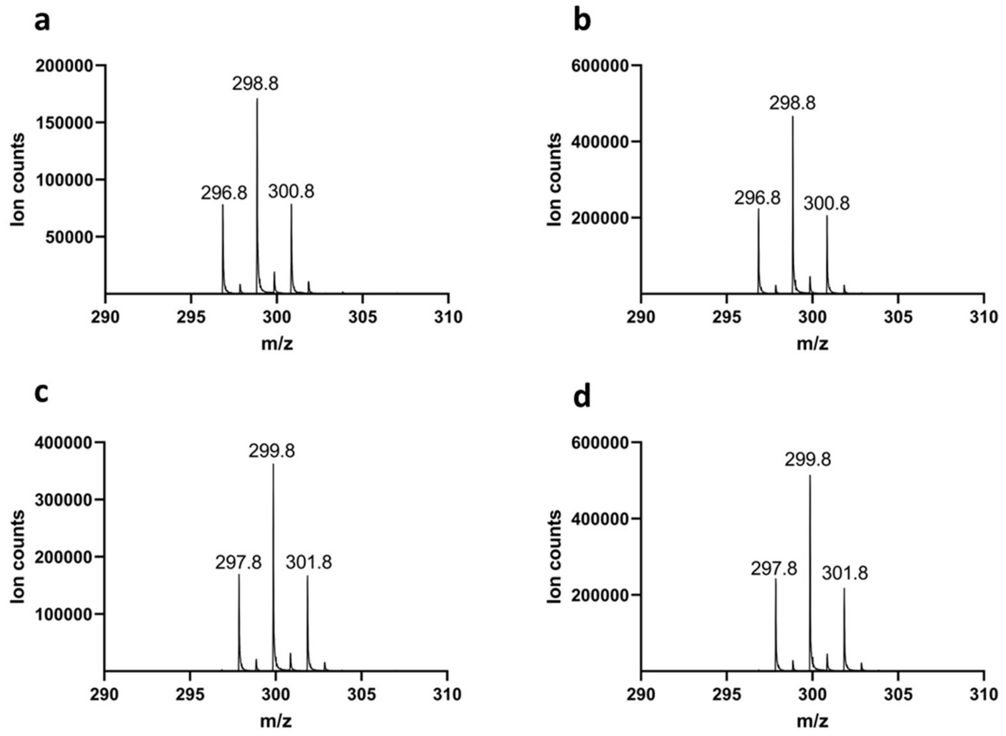

**Fig. S7 Mass spectra of the nitrile product when AetD reaction was performed with $^{13}$C-labeled isotopologues of substrate. a,** AetD + 5,7-dibromo-L-tryptophan. **b,** AetD + [2-$^{13}C_1$]-5,7-dibromo-L-tryptophan. **c,** AetD + [3-$^{13}C_1$]-5,7-dibromo-L-tryptophan. **d,** AetD + [1,2,3-$^{13}C_3$]-5,7-dibromo-L-tryptophan. A 1 Da increase in the mass of the nitrile product was observed only when [3-$^{13}C_1$]-5,7-dibromo-L-tryptophan and [1,2,3-$^{13}C_3$]-5,7-dibromo-L-tryptophan were used as substrates, confirming the $\beta$-carbon C-3 is retained in the product.

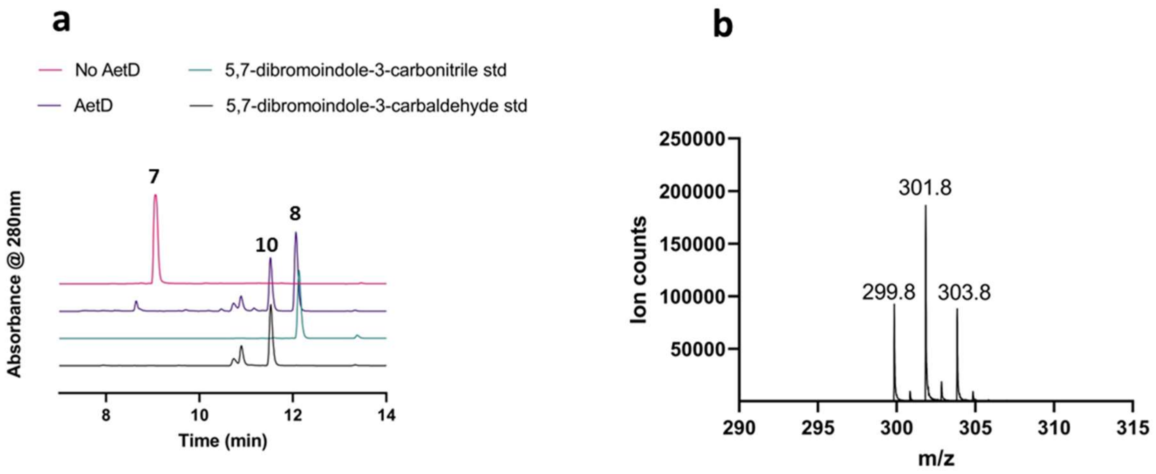

**Fig. S8 Characterization of the aldehyde shunt product formed in AetD reaction. a,** HPLC chromatogram showing the formation of 5,7-dibromo-indole-3-carbaldehyde (**10**) product in addition to the native 5,7-dibromo indole-3-carbonitrile (**8**) product. The retention time of the aldehyde shunt product formed in the AetD catalyzed reaction matches with that of an authentic standard. **b,** The observed mass of 5,7-dibromo-indole-3-carbaldehyde product in negative ionization mode.

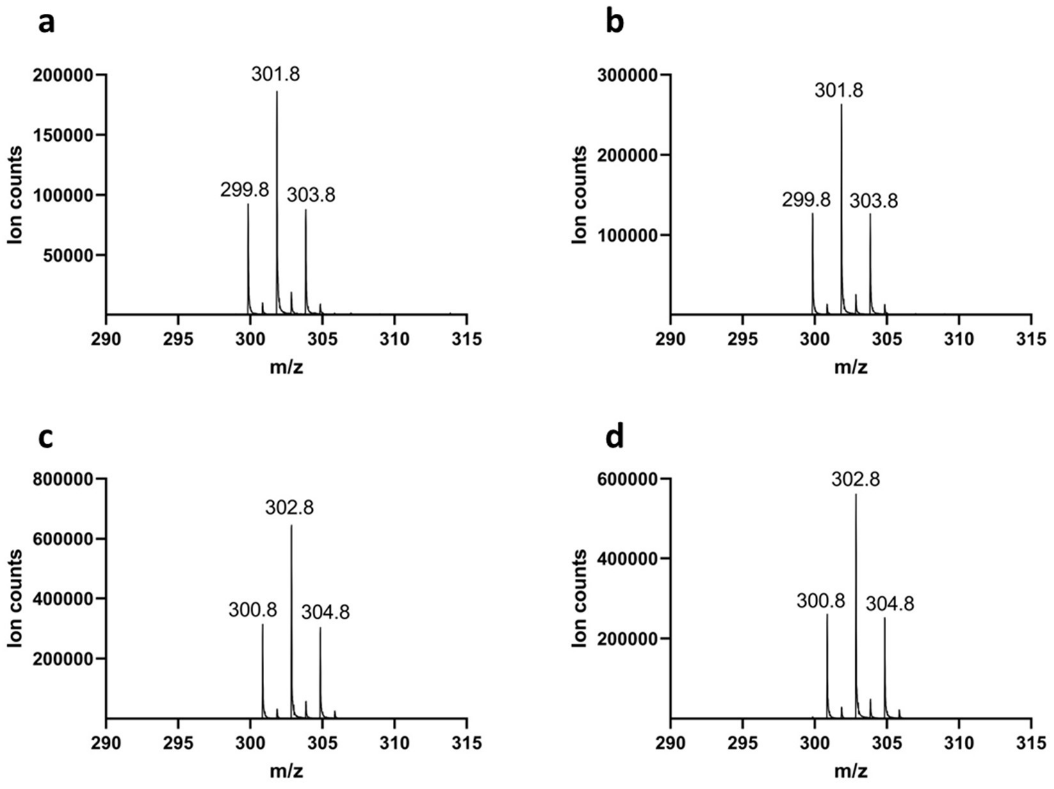

**Fig. S9 Mass spectra of the aldehyde shunt product when AetD reaction was performed with $^{13}C$-labeled isotopologues of 5,7-dibromo-L-tryptophan. a,** AetD + 5,7-dibromo-L-tryptophan. **b,** AetD + [2-$^{13}C_1$]-5,7-dibromo-L-tryptophan. **c,** AetD + [3-$^{13}C_1$]-5,7-dibromo-L-tryptophan. **d,** AetD + [1,2,3-$^{13}C_3$]-5,7-dibromo-L-tryptophan. A 1 Da increase in the mass of the aldehyde shunt product was observed only when [3-$^{13}C_1$]-5,7-dibromo-L-tryptophan and [1,2,3-$^{13}C_3$]-5,7-dibromo-L-tryptophan were used as substrates, confirming the β-carbon C-3 is retained in the shunt product.

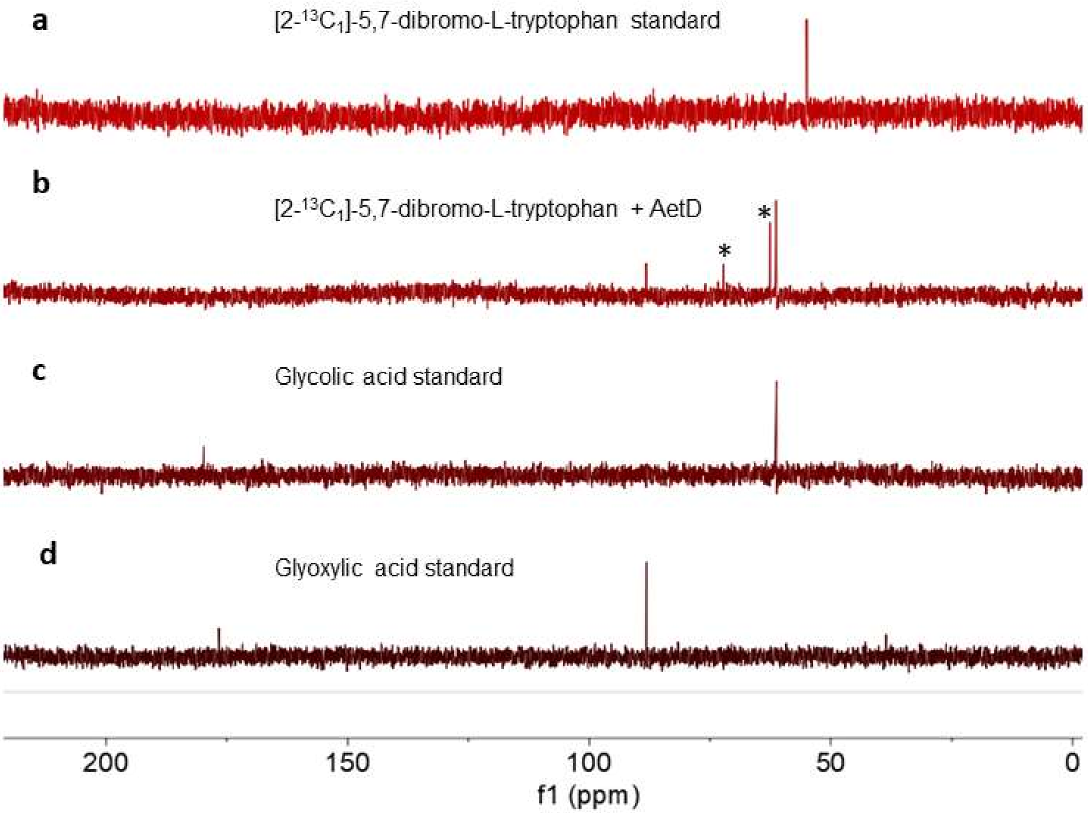

**Fig. S10 $^{13}$C NMR assay to identify the fate of carbon fragments.** Glycolic acid and glyoxylic acid were identified as the byproducts in the native and shunt pathways. **a,** [2-$^{13}C_1$]-5,7-dibromo-L-tryptophan standard. **b,** [2-$^{13}C_1$]-5,7-dibromo-L-tryptophan + AetD. **c,** Glycolic acid standard. **d,** Glyoxylic acid standard. When [2-$^{13}C_1$]-5,7-dibromo-L-tryptophan was incubated with AetD, two new NMR signals at 61 and 88 ppm appeared that match with glycolic acid (alcohol carbon) and glyoxylic acid (aldehyde carbon) standards, respectively. * Glycerol peak from the AetD protein stock.

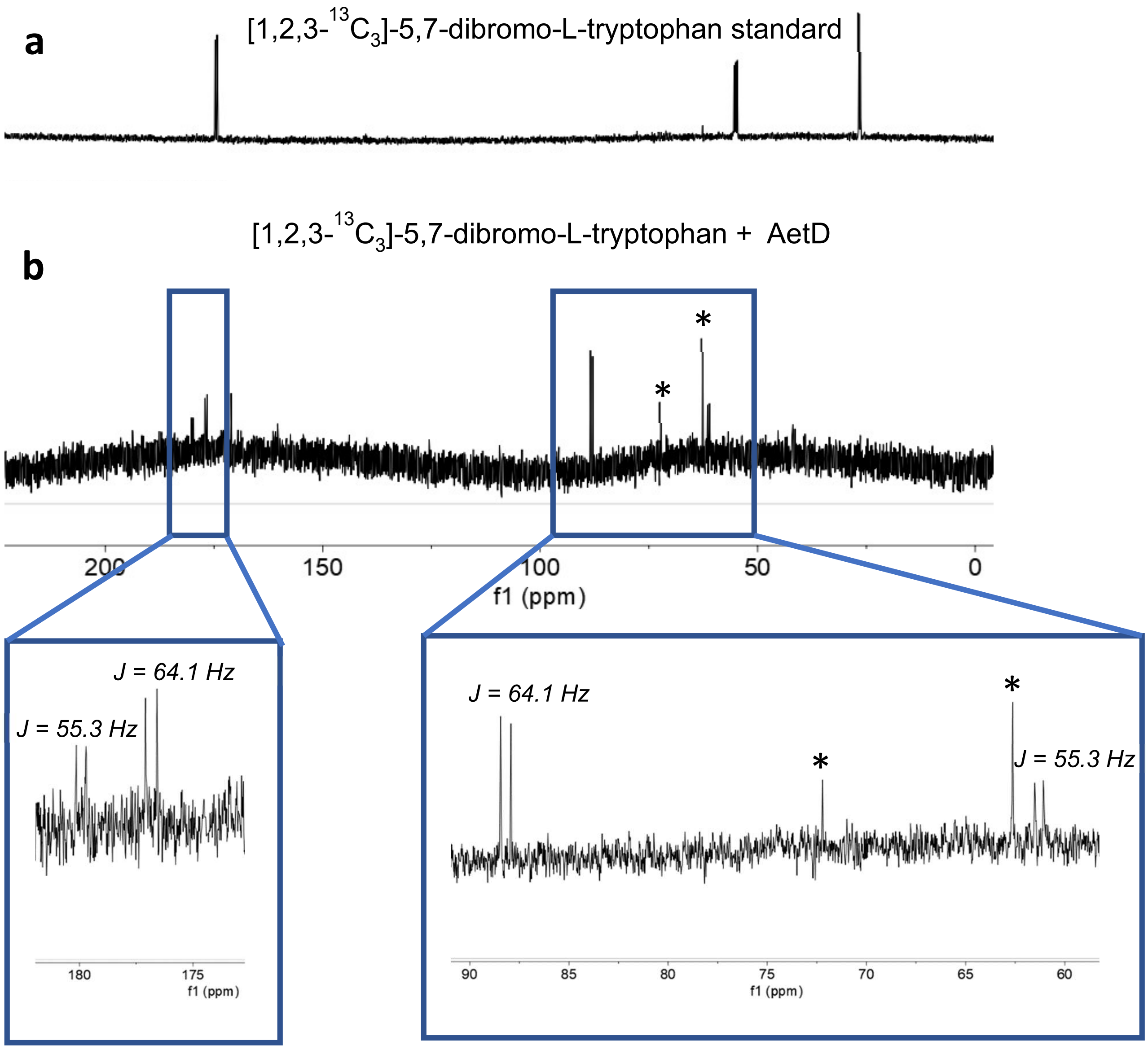

**Fig. S11 $^{13}$C-NMR assay with [1,2,3-$^{13}C_3$]-5,7-dibromo-L-tryptophan. a,** [1,2,3-$^{13}C_3$]-5,7-dibromo-L-tryptophan standard. **b,** [1,2,3-$^{13}C_3$]-5,7-dibromo-L-tryptophan + AetD enzyme assay. $^{13}$C-$^{13}$C NMR J-couplings were observed for both glycolic acid (55.3 Hz) and glyoxylic acid (64.1 Hz) generated in the AetD-catalyzed reaction. * Glycerol peaks from the AetD protein stock.

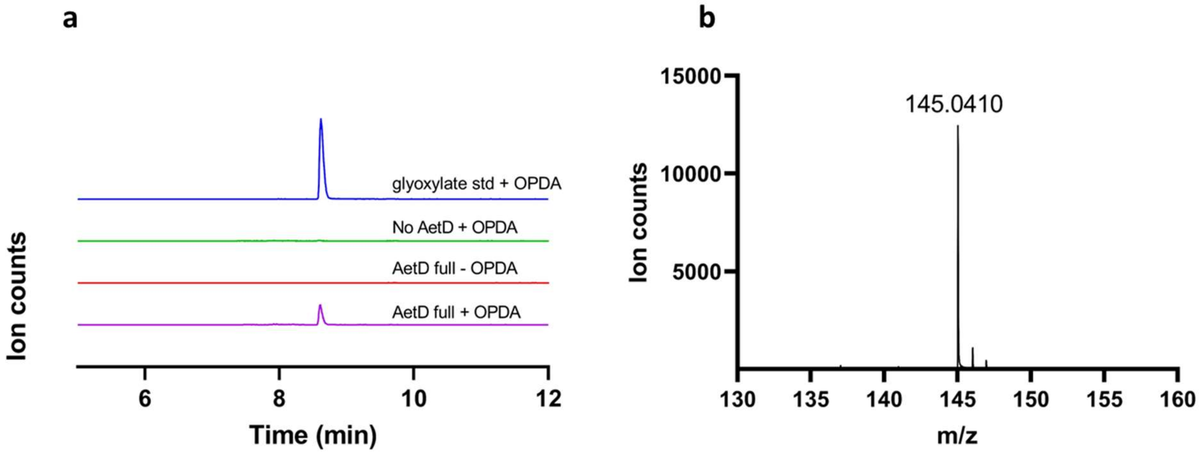

**Fig. S12 Detection of glyoxylate using *o*-phenylenediamine derivatization. a,** EICs at 145.04 corresponding to the *o*-phenylenediamine-glyoxylate $[M\text{-}H]^-$ adduct. The expected mass was only observed in the AetD full reaction in the presence of *o*-phenylenediamine. The mass cannot be detected in the absence of either AetD or *o*-phenylenediamine. **b,** The exact mass of *o*-phenylenediamine-glyoxylate adduct in negative ionization mode.

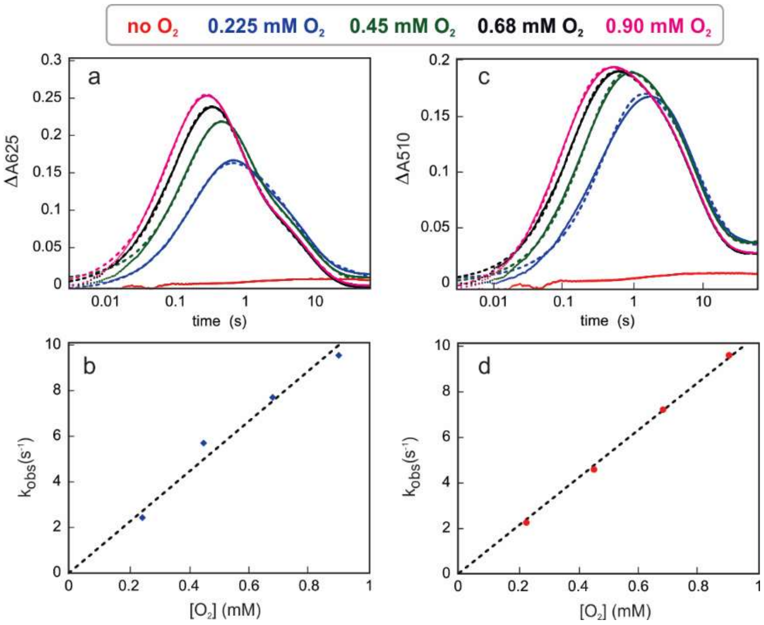

**Fig. S13** Kinetic traces showing the accumulation and decay of the absorbing intermediates as a function of time and $O_2$ concentration in the presence of 2 mM 5,7-dibromo-L-tryptophan. Experimental conditions: $O_2$-free solution AetD (0.30 mM), Fe(II) (0.60 mM, 2 molar eq), temperature: 5 ºC. Concentrations are reported prior to mixing (2-fold dilution).

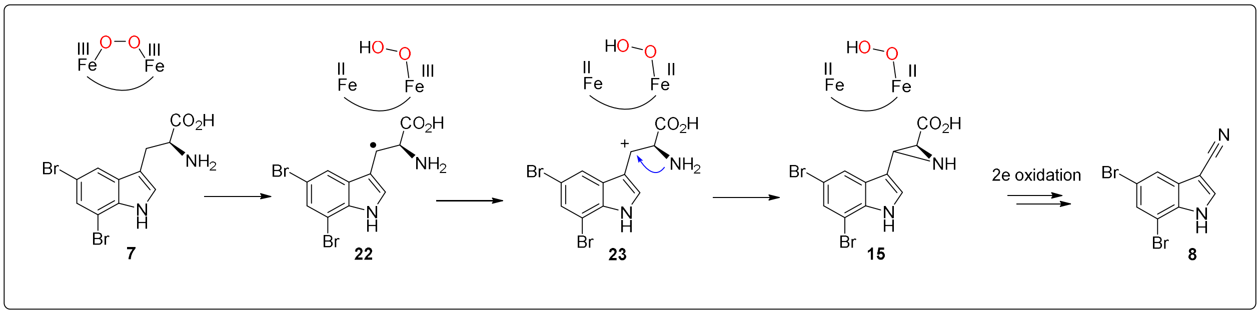

**Fig. S14 Carbocation-assisted mechanism of formation of the aziridine intermediate.**

**Table S1** AetD data collection and refinement statistics

| | Substrate-bound, Se-Met labeled | Substrate-bound | Substrate-bound and cofactor partially assembled | | Substrate-bound and cofactor fully assembled | |
|---|---|---|---|---|---|---|
| **Data collection** | | | | | | |
| Space group | $P\,2_12_12_1$ | $P\,6_522$ | $P\,2_12_12_1$ | | $P\,2_12_12_1$ | |
| Cell dimensions | | | | | | |
| *a*, *b*, *c* (Å) | 65.7, 85.8, 101.7 | 101.5, 101.5, 129.1 | 65.9, 86.1, 101.8 | | 65.9, 88.1, 102.8 | |
| α, β, γ (°) | 90, 90, 90 | 90, 90, 120 | 90, 90, 90 | | 90, 90, 90 | |
| | | | *Native* | *Fe peak** | *Native* | *Fe peak** |
| Beamline | SSRL 9-2* | APS 24-ID-C | SSRL 9-2 | APS 24-ID-C | APS 24-ID-C | |
| Wavelength (Å) | 0.97910 | 0.97918 | 0.95369 | 1.73404 | 0.97918 | 1.73404 |
| Resolution (Å) | 50.00-2.30 (2.44-2.30) | 50.00-2.08 (2.21-2.08) | 50.00-2.30 (2.44-2.30) | 50.00-2.29 (2.42-2.29) | 50.00-2.00 (2.13-2.00) | 50.00-2.49 (2.55-2.49) |
| Observed Reflections | 340402 (52130) | 205075 (30288) | 315639 (47689) | 189131 (30769) | 362648 (60171) | 246120 (9565) |
| Unique Reflections | 47930 (7654) | 22444 (3470) | 25947 (3958) | 48395 (7823) | 40402 (6430) | 40417 (2907) |
| $R_{sym}$ or $R_{merge}$ | 0.066 (0.304) | 0.131 (0.752) | 0.126 (0.626) | 0.124 (0.660) | 0.109 (0.543) | 0.162 (0.554) |
| $I / \sigma I$ | 22.8 (6.1) | 9.6 (1.6) | 15.6 (4.0) | 8.4 (2.2) | 13.19 (4.2) | 8.6 (3.1) |
| $CC_{1/2}$ (%) | 0.999 (0.967) | 0.995 (0.844) | 0.998 (0.927) | 0.993 (0.808) | 0.997 (0.937) | 0.987 (0.839) |
| Completeness (%) | 96.7 (95.5) | 92.4 (90.4) | 97.8 (93.6) | 95.0 (95.0) | 98.7 (99.0) | 99.4 (96.0) |
| Redundancy | 7.1 (6.8) | 9.1 (8.7) | 12.2 (12.0) | 3.9 (3.9) | 9.0 (9.4) | 6.1 (3.3) |
| **Refinement** | | | | | | |
| Resolution (Å) | | 43.97-2.08 | 39.53-2.30 | | 44.40-2.00 | |
| No. reflections | | 22435 | 25940 | | 40398 | |
| No. reflections $R_{free}$ | | 1122 | 1303 | | 2022 | |
| $R_{work}$ / $R_{free}$ | | 0.194/0.216 | 0.186/0.218 | | 0.184/0.209 | |
| No. atoms/molecules | | | | | | |
| Protein chain per ASU | | 1 | 2 | | 2 | |
| Protein atom | | 1985 | 3936 | | 3985 | |
| Protein residue | | 238 | 473 | | 480 | |
| Iron ion | | 0 | 3 | | 5 | |
| Substrate molecule | | 1 | 2 | | 2 | |
| Succinate ion | | 0 | 1 | | 0 | |
| Malate ion | | 1 | 2 | | 0 | |
| Glycerol molecule | | 0 | 1 | | 0 | |
| Water molecule | | 99 | 169 | | 261 | |
| *B*-factors (Å$^2$) | | | | | | |
| Protein Chains | | | | | | |
| Chain A | | 46.25 | 34.47 | | 32.03 | |
| Chain B | | - | 36.82 | | 32.43 | |
| Iron ion | | - | 45.45 | | 35.68 | |
| Substrate molecule | | 44.43 | 28.48 | | 26.75 | |
| Succinate ion | | - | 55.04 | | - | |
| Malate ion | | 52.75 | 33.59 | | - | |
| Glycerol molecule | | - | 41.94 | | - | |
| Water molecule | | 51.17 | 35.94 | | 36.77 | |
| R.m.s deviations | | | | | | |
| Bond lengths (Å) | | 0.002 | 0.002 | | 0.004 | |
| Bond angles (°) | | 0.59 | 0.58 | | 0.73 | |
| Ramachandran plot | | | | | | |
| Favored (%) | | 99.15 | 98.72 | | 98.95 | |
| Allowed (%) | | 0.85 | 1.28 | | 1.05 | |
| Outliers (%) | | 0.00 | 0.00 | | 0.00 | |

| | | | |
|---|---|---|---|
| Rotamer outliers (%) | 0.46 | 0.70 | 0.69 |
| Clashscore | 2.51 | 3.15 | 3.02 |
| No. TLS groups | 1 | 2 | 2 |

Values in parentheses are for highest-resolution shell. A single crystal was used to collect each dataset. *Statistics are for data processed anomalously (Friedel pairs scaled separately and unmerged).

### References

1 Adak, S., Lukowski, A. L., Schäfer, R. J. B. & Moore, B. S. From Tryptophan to Toxin: Nature's Convergent Biosynthetic Strategy to Aetokthonotoxin. *J. Am. Chem. Soc.* **144**, 2861-2866, (2022).

2 Bhandari, D. M., Fedoseyenko, D. & Begley, T. P. Mechanistic Studies on Tryptophan Lyase (NosL): Identification of Cyanide as a Reaction Product. *J. Am. Chem. Soc.* **140**, 542-545, (2018).

3 Kabsch, W. xds. *Acta Crystallogr. D* **66**, 125-132 (2010).

4 Karplus, P. A. & Diederichs, K. Assessing and maximizing data quality in macromolecular crystallography. *Curr. Opin. Struc. Biol.* **34**, 60-68 (2015).

5 Grosse-Kunstleve, R. W. & Adams, P. D. Substructure search procedures for macromolecular structures. *Acta Crystallogr. D* **59**, 1966-1973 (2003).

6 McCoy, A. J., Storoni, L. C. & Read, R. J. Simple algorithm for a maximum-likelihood SAD function. *Acta Crystallogr. D* **60**, 1220-1228 (2004).

7 McCoy, A. J. *et al.* Phaser crystallographic software. *J. Appl. Crystallogr.* **40**, 658-674 (2007).

8 Terwilliger, T. C. Maximum-likelihood density modification. *Acta Crystallogr. D* **56**, 965-972 (2000).

9 Jumper, J. *et al.* Highly accurate protein structure prediction with AlphaFold. *Nature* **596**, 583-589 (2021).

10 Emsley, P., Lohkamp, B., Scott, W. G. & Cowtan, K. Features and development of Coot. *Acta Crystallogr. D* **66**, 486-501 (2010).

11 Moriarty, N. W., Grosse-Kunstleve, R. W. & Adams, P. D. electronic Ligand Builder and Optimization Workbench (eLBOW): a tool for ligand coordinate and restraint generation. *Acta Crystallogr. D* **65**, 1074-1080 (2009).

12 Terwilliger, T. C. *et al.* Iterative-build OMIT maps: map improvement by iterative model building and refinement without model bias. *Acta Crystallogr. D* **64**, 515-524 (2008).

13 Morin, A. *et al.* Collaboration gets the most out of software. *elife* **2**, e01456 (2013).

14 Sugishima, M. *et al.* Crystal structure of dimeric heme oxygenase-2 from Synechocystis sp. PCC 6803 in complex with heme. *Biochemistry* **44**, 4257-4266 (2005).

15 Schwarzenbacher, R. *et al.* Structure of the Chlamydia Protein CADD Reveals a Redox Enzyme That Modulates Host Cell Apoptosis*. *J. Biol. Chem.* **279**, 29320-29324, (2004).

16 Zhang, B. *et al.* Substrate-Triggered Formation of a Peroxo-Fe2(III/III) Intermediate during Fatty Acid Decarboxylation by UndA. *J. Am. Chem. Soc.* **141**, 14510-14514, (2019).

17 McBride, M. J. *et al.* Substrate-Triggered µ-Peroxodiiron(III) Intermediate in the 4-Chloro-l-Lysine-Fragmenting Heme-Oxygenase-like Diiron Oxidase (HDO) BesC: Substrate Dissociation from, and C4 Targeting by, the Intermediate. *Biochemistry* **61**, 689-702, (2022).

18 McBride, M. J. *et al.* Structure and assembly of the diiron cofactor in the heme-oxygenase–like domain of the N-nitrosourea–producing enzyme SznF. *Proc. Nat. Acad. Sci. U.S.A* **118**, e2015931118, (2021).

19 Ng, T. L., Rohac, R., Mitchell, A. J., Boal, A. K. & Balskus, E. P. An N-nitrosating metalloenzyme constructs the pharmacophore of streptozotocin. *Nature* **566**, 94-99, (2019).
